# Colony-like Protocell Superstructures

**DOI:** 10.1101/2021.09.16.460583

**Authors:** Karolina Spustova, Chinmay Katke, Esteban Pedrueza Villalmanzo, Ruslan Ryskulov, C. Nadir Kaplan, Irep Gözen

## Abstract

We report the formation, growth, and dynamics of model protocell superstructures on solid surfaces, resembling single cell colonies. These structures, consisting of several layers of lipidic compartments enveloped in a dome-shaped outer lipid bilayer, emerged as a result of spontaneous shape transformation of lipid agglomerates deposited on thin film aluminum surfaces. Collective protocell structures were observed to be mechanically more stable compared to isolated spherical compartments. We show that the model colonies encapsulate DNA and accommodate non-enzymatic, strand displacement DNA reactions. The membrane envelope is able to disassemble and expose individual daughter protocells, which can migrate and attach via nano-tethers to distant surface locations, while maintaining their encapsulated contents. Some colonies feature ‘exo-compartments’, which spontaneously extend out of the enveloping bilayer, internalize DNA, and merge again with the superstructure. A continuum elastohydrodynamic theory that we developed reveals that the subcompartment formation must be governed by attractive van der Waals (vdW) interactions between the membrane and surface. The balance between membrane bending and vdW interactions yields a critical length scale of 273 nm, above which the membrane invaginations can form subcompartments. The findings support our hypotheses that in extension of the ‘lipid world hypothesis’, protocells may have existed in the form of colonies, potentially benefiting from the increased mechanical stability provided by a superstructure.

## Introduction

How the first biological cells came into existence on the early Earth is still an unanswered question. Significant research efforts have been directed towards the fabrication and studies of artificial compartments made from prebiotically plausible materials, i.e., the protocells, to program and understand their behavior and re-create the potential steps of development towards life. Since the first living cells are hypothesized to have a bilayer boundary^1^, and their modern counterparts also feature an enveloping bilayer, most protocell models are mammalian cell-sized spherical lipid-enveloped compartments, i.e. giant vesicles^2, 3^.

An individual protocell in an aqueous environment would have its membrane directly exposed to the fluid medium around it, subject to a plethora of mechanical, chemical and physicochemical challenges, such as shear stress due to agitation by currents and convection, osmotic stress, and pH fluctuations. An interesting alternative scenario was proposed to be the collective evolution of protocells, in the form of ‘vesicle colonies’^4^. Protocell colonies could have provided mutual benefits such as increased stability under harsh environmental conditions, and protocell communication by facilitating material exchange among the closely spaced compartments within the colony. High mechanical stability^5^ and enhanced material transfer would provide an advantage for primitive cells in the light of Darwinian evolution.

Agglomerates of giant unilamellar vesicles have been previously experimentally prepared, e.g. by bringing together initially isolated vesicles using adhesive molecules such as poly-L-arginine^6^, complementary DNA strands^7^, streptavidin-biotin^7^, divalent cations^8^ or oppositely charged biopolymers^9^. Similar structures were formed by directed assembly using instrumental techniques, including acoustical trapping^10^, optical tweezers^11^, and magnetic manipulation^12^. One example of a colony-like vesicular structure under prebiologically plausible conditions was reported, where osmotic shock of vesicles led to shrinkage of a lipid vesicle and formation of inverted daughter vesicles^13^. This transformation is presumably lipid-dependent, as vesicles made from certain lipid species did not exhibit this behavior^13^. Most recently, we showed the formation of colony-like model protocells emerging from the molecular lipid films on early Earth minerals and a Martian meteorite^14^.

Here we report the stepwise formation and growth of protocell superstructures containing tens to thousands of membranous compartments, originating from a single onion-shell lipid reservoir. Inside a lipid double bilayer compartment, which can be seen as a flat giant unilamellar vesicle (FGUV) spontaneously emerging from the reservoir by wetting a high energy surface, several layers of smaller vesicles grow from the surface up, leading to a densely packed pool of compartments of nearly identical shape and size, reminiscent of bacterial colonies. We predicted that the intra-vesicular compartments can grow from membrane undulations above a critical wavelength of 273 nm when the van der Waals interactions are attractive between the solid surface and the closely positioned membrane to it. The formed compartments then expand and fuse over time. Eventually, the original protocell, which serves as the enveloping layer and boundary for the colony, can disintegrate. We demonstrate that the protocell colonies, either enveloped in a bilayer or directly exposed, are mechanically more robust compared to individual compartments, can encapsulate DNA, and accommodate non-enzymatic strand displacement DNA reactions. Some colonies employ ‘exo-compartments’, which spontaneously extend out of the enveloping bilayer, uptake DNA and fuse again with the superstructure under re-distribution of its DNA content. The DNA-encapsulating colonies can disassemble, exposing individual daughter protocells that migrate and attach to distant surface locations via nano-tethers, while preserving their DNA content.

We report for the first time that distinctly substructured membrane containers with key features of protocells can autonomously and consistently emanate from lipid sources under conceptually simple, prebiotically relevant conditions. Our findings that the self-driven harvesting of surface free energy from solid surfaces for shape transformations introduce a new potential key step in the transition pathway from the non-living to the living world. We provide evidence that the evolution of organelles in primitive single cellular organisms does not necessarily require living cells as foundation, but can occur concomitantly with the self-assembly of single shell membranous structures from lipid material. The assembly route to physically separated, confined volumes in a protocell superstructure that can form colonies of enhanced mechanical stability, encapsulate molecules and pseudo-divide, creates new opportunities for prebiotic chemistry, with fresh options to overcome the limitations of “one-pot” syntheses that have thus far been explored. The potential of surface-free energy utilization for sophisticated self-organization processes by prebiotic assemblies has been overlooked in the past, a problem which we have addressed in this work.

## Results and Discussion

### Formation of superstructures

We experimentally observe the formation of up to several thousand micrometer-sized subcompartments inside a single unilamellar mother vesicle (**Fig. 1**). Prior to the formation process shown in **Fig. 1**, the unilamellar compartments were formed from a deposit of lipid material on solid substrates submerged in an aqueous environment (*cf*. **S1** for all utilized lipid compositions and type of surfaces). Ca^2+^ ions in the aqueous medium connect the lipid head groups with the solid substrate, and thereby establish pinning points at the solid-membrane interface, which promote adhesion of the mother vesicle to the solid surface^15–17^. **Fig. 1** shows the morphogenesis of the surface-adhered compartments after the adhesion was reversed: the surrounding Ca^2+^-buffer was gently exchanged with a Ca^2+^-free equivalent containing metal chelator molecules instead. The chelator forms a highly stable complex with Ca^2+^ and removes the pinning points from the nanogap between the membrane and the solid surface. This causes gradual de-wetting of the surface, and spontaneous formation of subcompartments^17, 18^. **Fig. 1a–d** show confocal micrographs of the subcompartmentalization over time, and **Fig. 1e–g** the corresponding schematic drawings. We recently reported on a related phenomenon, where unilamellar vesicles with no access to excess lipid material constituted model compartments, leading to the formation of subcompartments spread out on the basal membrane of the parent vesicle^18^. In the current study, each model mother protocell is physically connected to a multilamellar lipid reservoir (**Fig. 1a, e**). The reservoirs act as a material source, thus enabling the formation of densely packed layers of vesicular subcompartments inside the original compartment (**Fig. 1d, g**), and alleviate membrane tension upon external stress factors. Over the course of a few hours, almost the entire volume of the initial model protocell is filled from the surface up (**Fig. 1a–g**).

**Figure 1.**
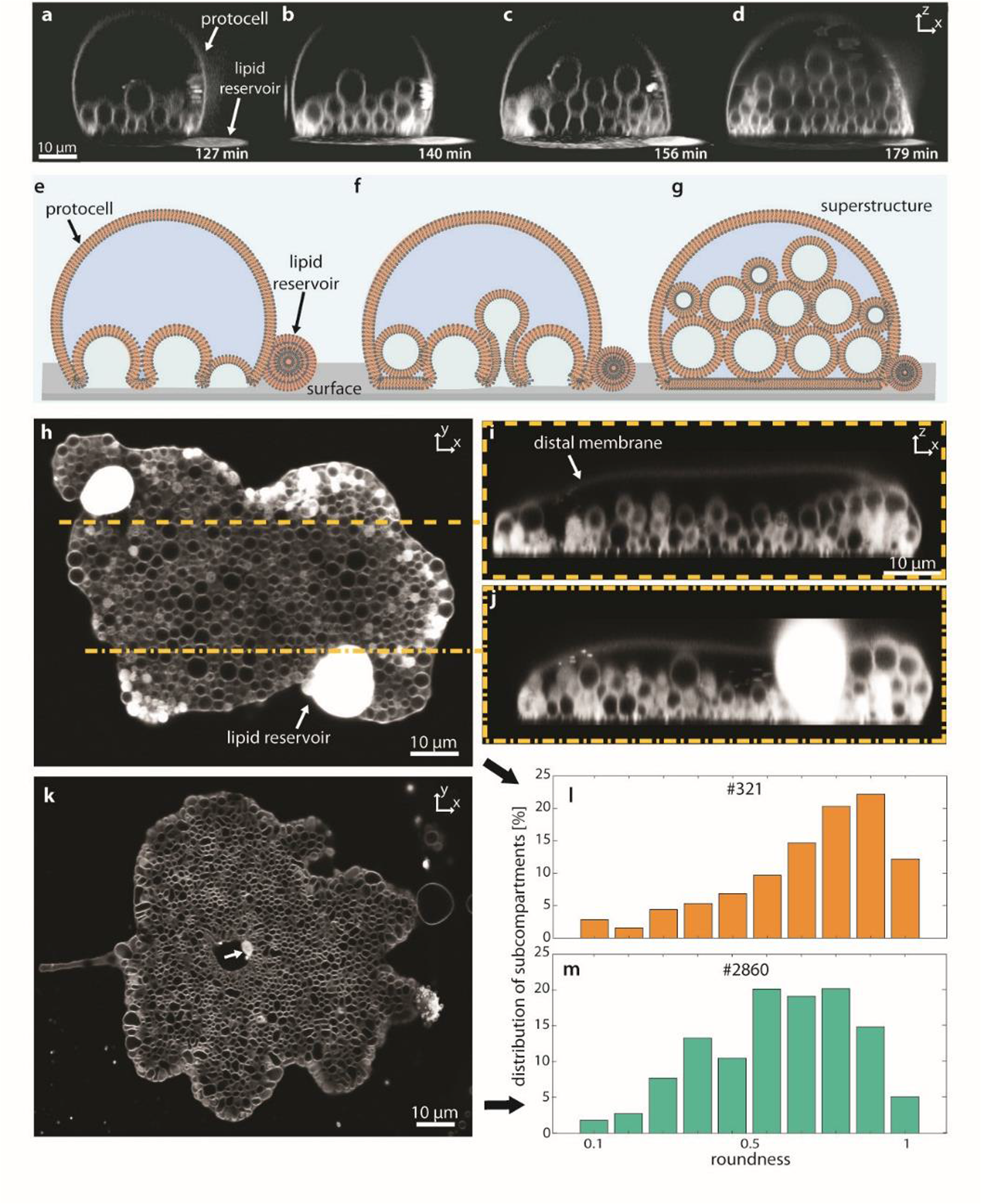
Formation of protocell superstructures. (**a-d**) Confocal micrographs showing the gradual formation of subcompartments over time (x-z cross sectional view). (**e-g**) Schematic drawings corresponding to (**a-d**). The protocell enveloping the subcompartments is connected to a multilamellar lipid reservoir. Over time, multiple layers of spherical subcompartments form inside the protocell from the bottom (solid surface) up. Eventually the compartments fill up the original protocell volume (**d** and **g**). (**h, k**) Confocal micrographs of large protocell superstructures (x-y cross sectional view). (**i-j**) shows the x-z cross sectional view along the dashed lines in (**h**). Up to 3 layers of subcompartments were observed in this experiment. (**l-m**) Roundness analyses of the superstructures shown in (**h**) and (**k**), respectively. The structures in (**i**) have subcompartments with higher roundness (closer to 1). The number of compartments in (**h**) is 321 and in (**k**), 2860. In (**k**) the multilamellar reservoir is mostly consumed (white arrow).

The mother protocell spreads on the solid substrate before de-wetting occurs, drawing lipid material from the multilamellar reservoir. Thus, the protocell membrane can expand across the *xy* plane over a wide area of several hundreds to thousands of square micrometers in the form of a flat unilamellar vesicle^19^. This initial spreading is followed by the subsequent steps of chelation, de-wetting and formation of subcompartments. **Fig. 1h–k** contain two examples of resulting structures. **Fig. 1i–j** show the cross-sectional profile views along the dashed lines in **Fig. 1h**. Multiple layers of subcompartments can be observed within the protocell internal volume, similar to the structure in **Fig. 1d**, just extended over a wider surface area. **Fig. 1l–m** depicts the distribution of roundness values^20, 21^ of the individual compartments in **Fig. 1h** and **k**. The compartments in panel (**h**) adopt more round shapes (perfectly round:1, highly elongated: 0.1) compared to the ones in (**k**). If the initial spreading is prevented due to surface impurities, roughness or pinning, the subcompartments fill inside a somewhat constrained volume (**Fig. 1k**). This will result in the mild shape deformations of the internalized compartments leading to the reduced roundness values.

We term these lipid architectures, which emerge from the surface up, ‘protocell superstructures’ (**Fig. 1g**). Superstructures contain multilayers of densely packed membranous compartments of identical shape and size, all originating from the same membrane assembly autonomously, resembling colonies of single cell organisms^22^. To this respect, ‘vesicle colonies’ in the context of the origins of life have been discussed earlier as a possible step paving the way for the first biological cells^4^.

Our experimental system consists of a minimal number of prebiotically relevant components: amphiphiles, a solid surface, and an aqueous environment suitable for surfactant self-assembly. Our findings presume that phospholipids, variants of which we have used for the experiments, could have been present under prebiotic conditions^23–26^. Adhesion promoting Ca^2+^ ions were present in early Earth minerals, which could have acted as a source in a natural prebiotic environment^27, 28^. The chelator molecules EDTA and BAPTA used in our experiments to induce de-pinning were highly likely not present on the early Earth. However, clay minerals which are known to adsorb mono- or divalent cations^29^ could have potentially acted as sources for chelating agents. We consider experimental studies to this respect a valuable next step in order to test this hypothesis.

The subcompartments are held together by the enveloping membrane of the original model protocell, which creates a physical super-boundary. This outer membrane shell can, however, disintegrate to free the subcompartments as independent daughter cells^18^, which maintain their connection to the solid surface while remaining in close proximity to each other (*cf.* **SI Fig. S2**).

### Growth and dynamics of superstructures

Growth and merging of small subcompartment colonies inside the flat giant unilamellar model protocell compartments were followed by confocal microscopy (**Fig. 2, SI Movie S1**). **Fig. 2a–i** depicts an example in which more than 300 subcompartments form over the course of a few minutes (top view). Magnifications in **Fig. 2b, e, h** show the regions framed in dashed lines in **Fig. 2a, d, g**. Panel (**b**) shows merging of small colonies (>3 subcompartments), (**d**) the expansion of existing colonies due to the formation of new subcompartments, and (**h**) fusing subcompartments within the colonies. A mild temperature increase was applied during the experiment shown in **Fig. 2** in order to shorten the transformation time period of the experiment^18, 30^. **Fig. 2j** shows the average (arithmetic mean) area of the isolated colonies (orange plot) as well as the average area of each subcompartment (blue plot) in **Fig. 2a–i** over time. The graph in **Fig. 2k** shows the number of colonies (orange plot) and the number of individual subcompartments (blue plot). The gray zones in both graphs indicate the time period during which colonies merged and grew in size. Upon merging, the total number of colonies is reduced. The zone depicted in pink covers the period in which the subcompartments were fusing (growth in average size, drop in number), and the colonies were merging (number of colonies decreasing, and their size increasing). The total area occupied by the colonies was increasing over time (**Fig. 2l**). After the consumption of most of the lipid reservoir (arrow in **Fig. 2a**), it remained constant (> ~4 min.). The total perimeter of the colonies decreased over time (**Fig. 2m**). **Fig. 2l–m** shows that the superstructure minimizes its surface free energy over time by minimizing its surface area to perimeter ratio. **SI Movie S1** contains three examples of formation and merging of the colonies. The dynamic motion of clusters, i.e., merging and collective motility, resembling the behavior reported for bacteria populations^31, 32^. Another example, which we deem morphologically and dynamically similar, is surfactant foam^33, 34^.

**Figure 2.**
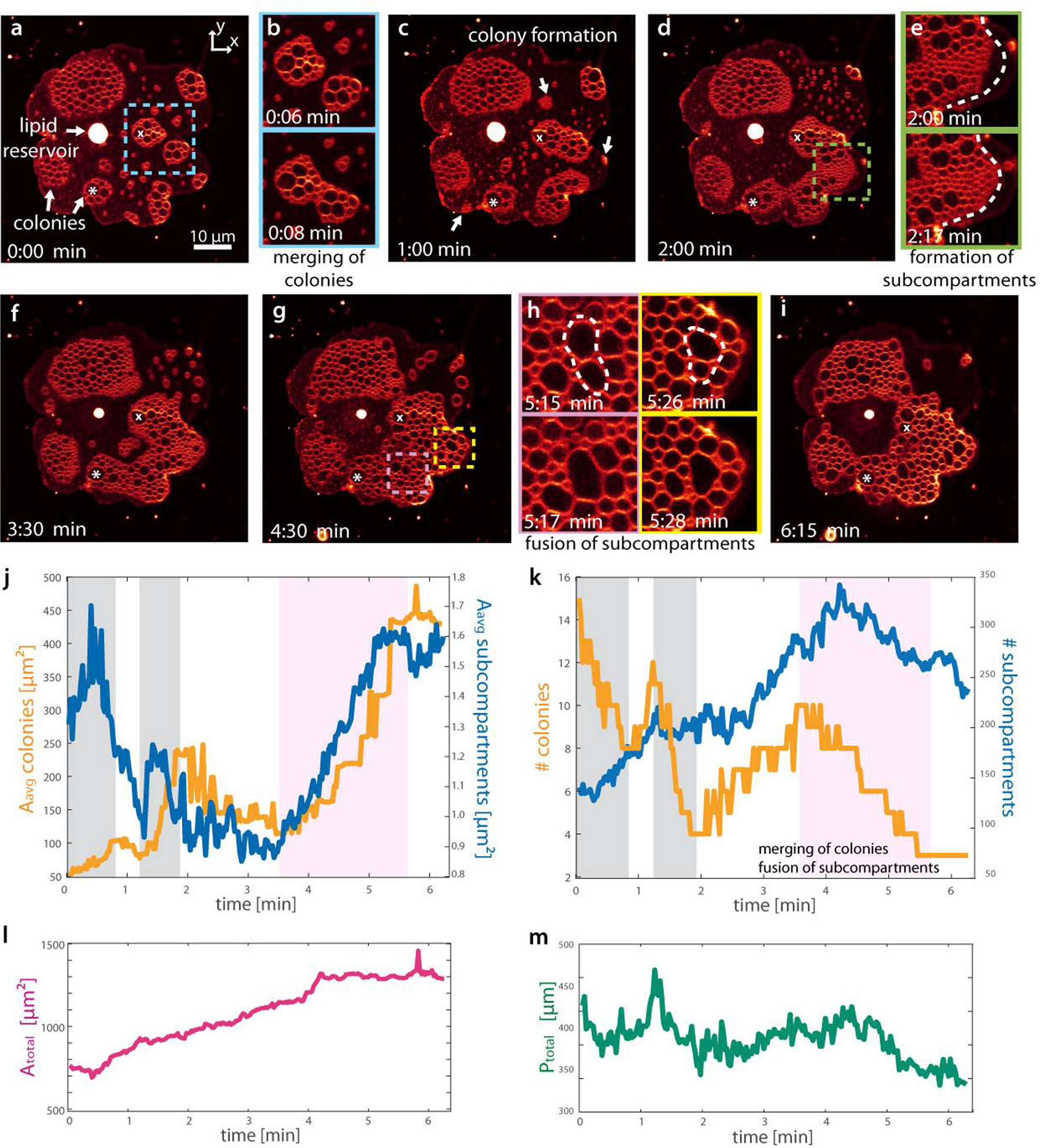
Growth and dynamics of protocell superstructures. (**a-i**) Confocal microscopy time series showing the emergence, growth and fusion of protocell superstructures (x-y cross sectional view). The regions framed in dashed lines in (**a, d**, **g**) are shown in (**b, e, h**), respectively. (**b**) Merging of two initially isolated colonies. (**e**) Emergence of new compartments leading to the expansion of a colony. (**h**) Fusion of two adjacent subcompartments. Two selected subcompartments, marked with ‘x’ and ‘*’, have been monitored throughout the experiment in (**a-i**). (**j**) Average area of colonies (orange plot) and average area of individual subcompartments (blue plot), vs. time. The gray zones (t < 3 min.) indicate that new subcompartments are continuously forming, leading to new colonies, their growth and merging. The pink zone (t> 4 min) indicates the period in which growth and expansion of existing clusters are observed. (**k**) Number of colonies (orange plot) and subcompartments (blue plot) vs. time. (**l**) Total area of the colonies vs. time. (**m**) Total perimeter of the colonies vs. time.

### Continuum theory for the onset of subcompartment formation

To explain the dynamics of the early subcompartment formation, we developed a continuum theory that couples the elasticity of membrane deformations with the induced viscous flow of the solvent underneath the membrane. When the proximal (surface-adhered) lipid bilayer is detached from the substrate due to removal of pinning points, the thermal energy *K*_*B*_ *T* (*K*_*B*_: Boltzmann’s constant, *T*: temperature) must result in fluctuations in the lipid bilayer membrane, causing it to lift off at a height *h* from the underlying charged substrate (**Fig. 3**). We considered that an elastohydrodynamic instability can amplify these fluctuations and lead to the onset of inward growth of subcompartments. The three potential causes for the instability must then be (i) electrostatic repulsion between the membrane and the substrate, (ii) a possible spontaneous curvature *c*_0_ of the membrane^35^, (iii) attractive van der Waals (vdW) interactions between the membrane and the substrate when the distance in-between is about the bilayer thickness (*h* ≲10 nm). The role of electrostatic repulsion can be omitted due to screening by the ionic ambient solution, since the bilayer thickness (≲10 nm) is much bigger than the Debye screening length of the solvent *λ*_*D*_=0.84 nm^36^. Furthermore, the lipid composition in each monolayer leaflet of the membrane is identical, therefore the spontaneous curvature *c*_0_ must be zero. Even if *c*_0_ were finite on a flat membrane, the energy gain, e.g. at a crest of a plane wave undulation with a curvature *c*_0_ would be balanced by the energy cost at a neighboring trough with the curvature −*c*_0_. Consequently, the wave-like undulation would altogether be suppressed by the membrane bending energy. On the other hand, in our system, the vdW interactions between the lipid bilayer and the aluminum substrate are attractive (set by the Hamaker constant *A*_*H*_≈2×10^−21^ J >0) and thus destabilize a free interface that confines a liquid^37, 38^: Since vdW interactions decay with increasing distance, any forming crest would be attracted to the substrate less than the neighboring troughs, creating a pressure gradient from the crests to troughs. This in turn leads to an influx of the aqueous ionic background solvent under the proximal membrane (characterized by the vesicle radius *L*≳1 μm, **Fig. 3a**), reinforcing the membrane fluctuations in a positive feedback cycle. The bilayer-solvent interfacial tension is *γ*≈0, thus, a perturbed membrane region could bulge indefinitely as soon as it acquires a spherical profile during growth since the Laplace pressure across the membrane bilayer would then vanish when *c*_0_=0.

**Figure 3.**
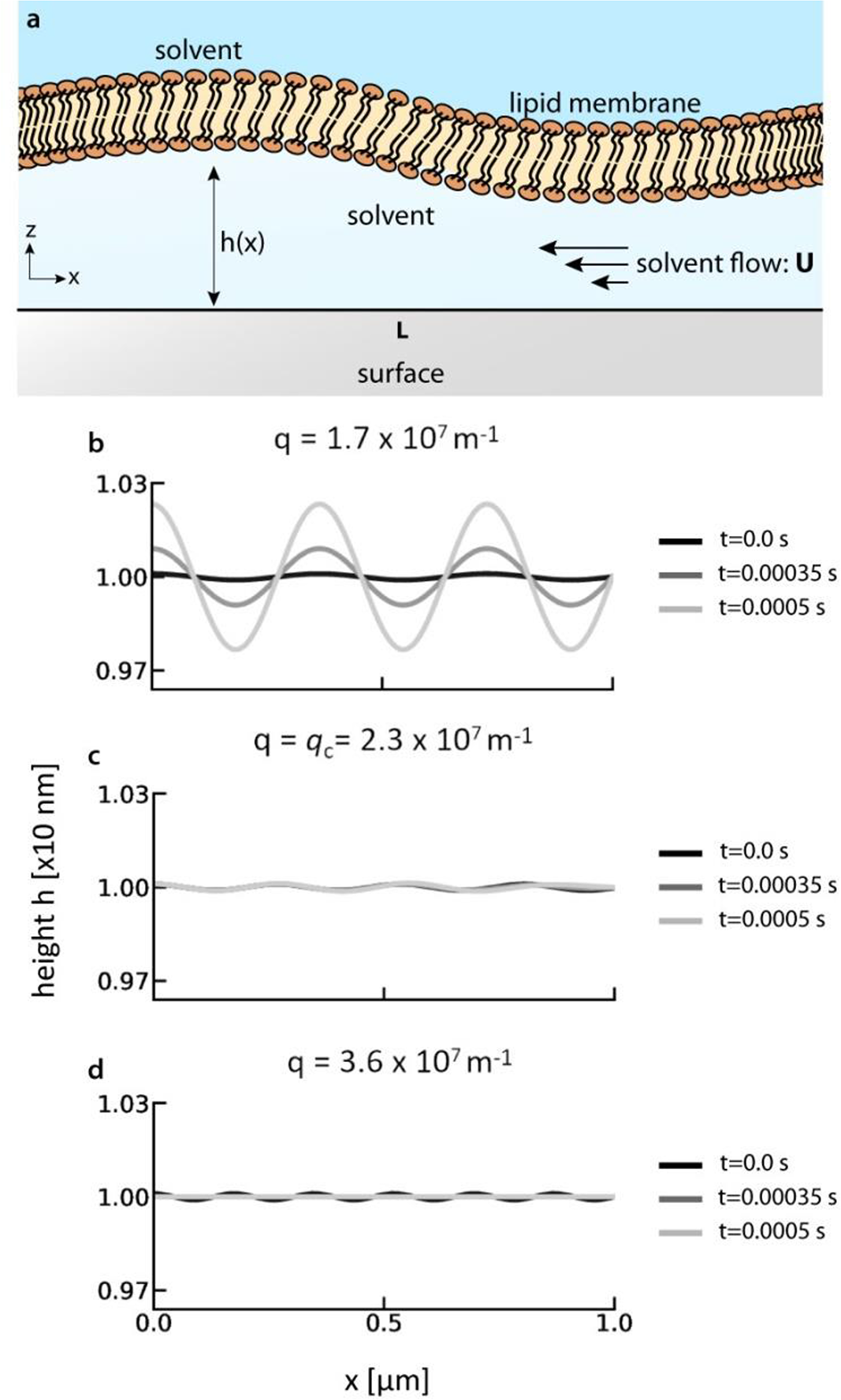
Elastohydrodynamic instabilities as a mechanism for subcompartment formation. (**a**) The bilayer-solvent (water)-substrate system. At the proximal membrane, the lipid bilayer and the solid surface confine a thin film of aqueous ionic solvent, which flows in response to the pressure gradients along the *x*−direction. The center point at the base of the vesicle corresponds to the left boundary, and the right boundary is located in the proximity to the vesicle front. (**b-d**) Numerical results for the height of the membrane in linear deformation regime. Height profiles for (**b**) *q* < *q*_*c*_, (**c**) *q* = *q*_*c*_ and (**d**) *q* = *q*_*c*_ at three different times. The center point at the base of the vesicle corresponds to *x*=0 μm, and the vesicle front is located at *x*=1 μm. The grayscale corresponds to three different times t=0 s, t=0.00035 s and t=0.0005 s, respectively. The results from numerical solutions show emergence of the instability below the critical wave number *q*_*c*_=2.3×10^7^ m^−1^, equivalently above critical wavelength *λ*_*c*_ = 2*π*/*q*_*c*_~273 nm.

To quantify the size of bulging at the proximal membrane of a lipid bilayer vesicle, we developed an elastohydrodynamic thin film theory that relates the membrane bending and the attractive vdW interactions to the pressure gradients driving solvent flow in a high-aspect-ratio domain (*h* ≪ *L*) (**Fig. 3a**). For simplicity, we took a two-dimensional (2D) cross-section of the proximal membrane and assumed that chelation occurs uniformly along the horizontal direction at time *t*=0 of the simulations. The small fluctuations are represented by a dynamic term *h*_1_(*x*, *t*) ≪ *h* that serves as a correction term to the equilibrium bilayer height *h*_0_>0, that is, *h* = *h*_0_ + *h*_1_. In the limit |*h*| ≪ *h*_0_ ≪ *L*, the mass conservation of the incompressible solvent-bilayer system and the flow induced by the pressure gradients due to membrane bending and vdW interactions yield the linear evolution equation:

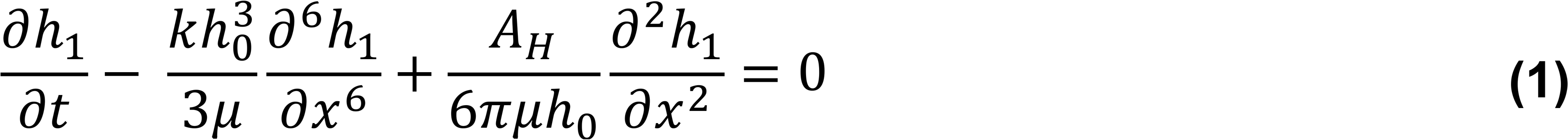

for the height fluctuation profile *h*_1_ where *k*~10^−19^ J is the membrane bending modulus, and μ~10^−3^ Pa·s is the dynamic viscosity of the aqueous solvent. The onset of bulging can be determined by the linear stability analysis of **Eq. 1** when the height undulations *h*_1_ represent plane wave deformations, such that *h*_1_ = |*h*_1_|*e*^(*iqx*+*st*)^, *q* being the wave number and *s* being the inverse time scale of fluctuations. Substituting *h*_1_ in **Eq. 1** returns the dispersion relation with a critical wave number *q*_*c*_

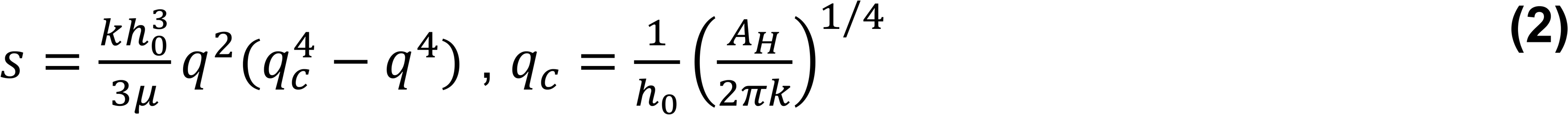

**Eq. 2** reveals that when *q* < *q*_*c*_, *s* > 0, i.e., the vdW interactions dominate the membrane bending, destabilizing the interface. Conversely, when *q* ≥ *q*_*c*_, *s* ≤ 0, the bending energy would suppress the membrane fluctuations against the attractive vdW interactions. Using the experimental values *h*_0_≈10 nm, *A*_*H*_≈2.08×10^−21^ J, and *k*≈1.2×10^−19^ J^39^ returns *q*_*c*_=2.3×10^7^ m^−1^, equivalently *λ*_*c*_ = 2*π*/*q*_*c*_ ~273 nm. This means that the size of the perturbed membrane regions larger than ~273 nm will lead to the growth of subcompartments. We cannot determine the exact size of the nucleating compartments with confocal microscopy, as compartments smaller than the resolution limit (150 nm) will not be distinctly visible, however, we can confirm that the smallest subcompartments we can observe are of submicron size (**Fig. 2d**).

To investigate the dynamics of the bilayer height profile *h* in the presence of attractive van der Waals interactions, we solved **Eq. 1** numerically in the stable regime *q* ≥ *q*_*c*_ and in the unstable regime *q* < *q*_*c*_. The simulation details, the initial condition for *h*_1_ determined by the thermal fluctuation spectra, and the physical boundary conditions for **Eq. 1** are summarized in the Methods. **Fig. 3** presents the time evolution of the membrane height *h* for the wave numbers *q*=1.7×10^7^ m^−1^ (**Fig. 3b**), *q* = *q*_*c*_=2.3×10^7^ m^−1^ (**Fig. 3c**), and *q*=3.6×10^7^ m^−1^ (**Fig. 3d**). Below the critical wave number *q* < *q*_*c*_, the inverse time scale *s* is positive such that fluctuations grow in time, leading to an instability (**Fig. 3b**). In this regime, the flow profile due to the pressure gradient further amplifies the height undulations (**Fig. S3.2g**). At the critical wave number *q* = *q*_*c*_ the height profile remains stationary since *s* vanishes (**Fig. 3c**, **Eq. 2**). This steady state is characterized by zero pressure and vanishing flow throughout the system (**Fig. S3.2e** and **h**). In **Fig. 3d**, *s* is negative (*q* > *q*_*c*_), and the fluctuations die out as the resulting flow suppresses the height undulations (**Fig. S3.2i**). These results suggest that the height instability triggered by the attractive van der Waals interactions between the lipid bilayer membrane and the aluminum substrate is a plausible mechanism for the nucleation of subcompartments. We also performed stability analyses for the screened electrostatic interactions and the effect of the spontaneous curvature. We validated that both effects stabilize the bilayer profile at a linear order (*cf.* **SI Section 3**).

### Mechanical stability of the superstructures

To characterize the durability of the protocell superstructures in alternating aqueous environments, we investigated the effect of different osmotic conditions on the protocell superstructures (**Fig. 4**). Isolated giant unilamellar vesicles (**Fig. 4a–b**), enveloped and non-enveloped vesicle colonies (**Fig. 4c–e**), were consecutively exposed to hypotonic (low salt) and hypertonic (high salt) solutions. **Fig. 4f** shows the duration for which each of the structures remains intact under the applied osmotic stress. For 30 s, all structures were initially exposed to a gentle hydrodynamic flow (10−100 nl/s) from an open-volume microfluidic device^40^. The lipid structures were then exposed to a hypotonic solution for 30 s, upon which all isolated lipid vesicles immediately ruptured (asterisks in panel **f**). The effect of imbalanced osmotic conditions on the deformation or rupture of individual compartments has been discussed previously^41^. Both the non-enveloped and enveloped structures maintained their integrity during this period. Subsequently, a hypertonic solution was applied, which rapidly caused the non-enveloped vesicle clusters (**Fig. 4c**) to de-cluster and some to disintegrate (**Fig. 4d–e**). The superstructures enveloped in a bilayer remained intact for another 2 minutes (*cf.* **SI Movie S2** for the entire period of the experiment and other examples).

**Figure 4.**
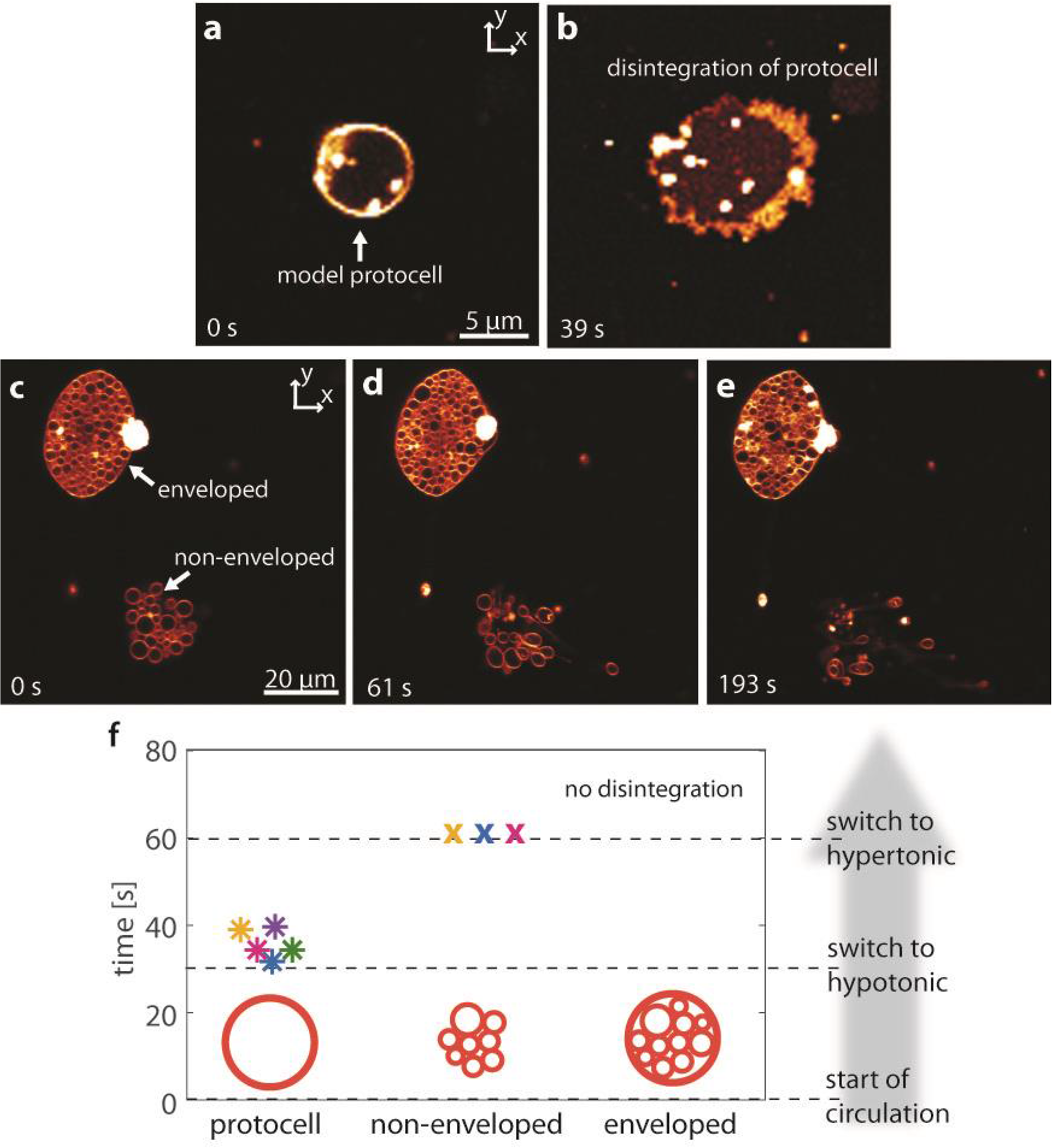
Mechanical stability of protocell superstructures. Confocal micrographs of (**a-b**) an isolated lipid compartment, (**c-e**) protocell superstructures with and without a membrane envelope. (**f**) Time periods of each type of model protocell structures in (**a-e**) during exposure to hypo- and hypertonic environments. The gray arrow shows the order of events along the y axis (time). At t=0 s the structures are exposed to a gentle hydrodynamic flow via an open volume microfluidic device. At t=30 s the structures are exposed to hypotonic solution, upon which the isolated protocells disintegrate (asterisks). At t=60 s the protocell superstructures are exposed to hypertonic solution. Shortly thereafter, the non-enveloped superstructures are de-assembled (crosses). The enveloped superstructures maintain their integrity as long as 210 seconds at which the experiment is terminated.

The results shown in **Fig. 4** support the earlier hypotheses that protocell colonies have a higher mechanical stability^42^. Even the non-enveloped structures withstand the applied ambient stress conditions better than the individual vesicles. This might be due to the presence of lipid nanotubes which tether each compartment to the surface. Membrane nanotubes can act as reservoirs to balance increase membrane tension and increase robustness of lipid compartments^43^. The enveloped superstructures exhibited higher mechanical stability compared to non-enveloped colonies upon hypertonic shock (**Fig. 4e**). This suggests that the enveloping membrane of the superstructure protects the colony from the initial osmotic shock by acting as a barrier, and reshapes itself in response^44^, possibly also relieving membrane tension by drawing additional lipid material from the attached multilamellar reservoir. It was shown recently that hypertonic conditions cause the compression of magnetically-assembled vesicle clusters and decrease the permeability into the cluster^12^. In contrast, vesicle colonies bridged by homopolyamino acids exhibited less physical stability than single compartments when exposed to osmotic stress, likely due to membrane perturbations caused by the anchoring of the amino acids^6^.

### Encapsulation of compounds

Chemical information exchange within cell colonies or across generations requires distinct compartments of reactants. We studied the ability of the superstructures to encapsulate compounds from the ambient environment, e.g. DNA (**Fig. 5, SI Movie S3**) and fluorescein (**SI Fig. S4**). Encapsulation of genetic fragments is considered an important step in the evolution of primitive cells^45^. A membrane-bound compartment provides confinement, and protection from parasitic genetic fragments, and can increase reaction rates, which could have greatly enhanced protocell evolution, e.g. in an RNA world^46, 47^. RNA encapsulation and distribution in similar lipid systems were recently reported^30^.

**Figure 5.**
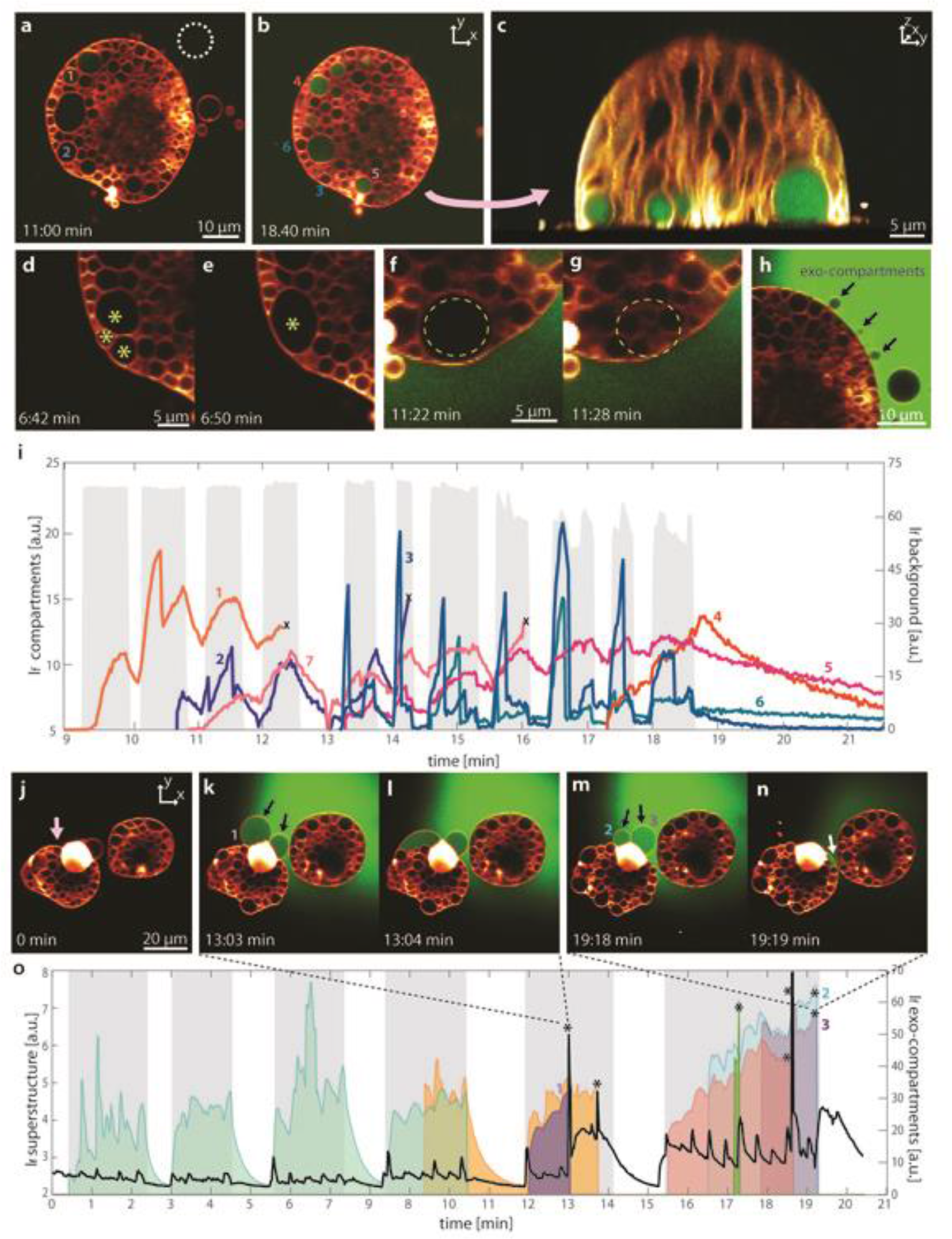
DNA encapsulation. **(a-h**) Confocal micrographs showing the fluorescently labeled DNA spontaneously encapsulated inside protocell superstructures. (**a-b**) Micrographs show a superstructure after 11 min (**a**), and after 18 min (**b**), of DNA exposure. (**c**) A confocal micrograph in y-z cross sectional view, showing the compartments encapsulating labeled DNA. Close up view of: (**d-e**) merging subcompartments, (**f-g**) a large subcompartment spontaneously substituted with multiple small subcompartments, (**h**) exo-compartments. (**i**) Graph showing the FAM-DNA fluorescence intensity over time starting from t=9 min of exposure to DNA. The width of the gray zones indicates the duration of DNA superfusion, and their height, the intensity of the DNA in the ambient solution (y axis on the right of the graph: background fluorescence I_f_). Each of the colored plots correspond to a subcompartment in (**a-b**). The number of each plot matches the associated compartment labeled with the same number in (**a-b**) (an exception is plot 7 which is not visible in (**a-b**)). Plots in the blue spectrum represent the subcompartments that take up the DNA and immediately lose it once superfusion is terminated. Plots in red spectrum represent subcompartments which maintain DNA even the superfusion is terminated. The plots marked with ‘x’ show subcompartments which spontaneously disappeared/relocated and could not be monitored further. (**j-n**) Confocal micrographs showing the fusion of exo-compartments to the protocell superstructure and redistribution of contents (arrow in **k, m, n**) (**o**) FAM-DNA fluorescence intensity of the superstructure marked with pink arrow in **(j)**, over time. The black plot shows the average FAM-DNA intensity inside the superstructure. The gray zones indicate DNA superfusion periods (green background in **j-n**). The areas under the plots have been colored to facilitate visualization. Each colored plot shows the FAM-DNA intensity of an exo-compartment vs. time. The time points at which the exo-compartments are merging with the protocell superstructure are marked with an asterisk.

**Fig. 5a–h** shows a confocal microscopy time series of a superstructure which can encapsulate fluorescently labeled single-stranded DNA (20 nucleobases). Panel **(c)** shows a 3D micrograph, whereas the other panels show a cross-sectional top view. We used the open volume microfluidic pipette described earlier^40^ to superfuse the superstructure locally with labeled ssDNA. The colony of subcompartments inside the superstructure was dynamic during the DNA exposure. In **Fig. 5d–e**, fusion of 3 adjacent subcompartments is shown (asterisks in **d-e**). In another case, one large, distinct compartment (**Fig. 5f**) was replaced by 5 smaller compartments (**Fig. 5g**). We also observed the formation of ‘exo-compartments’ which extended out of the protocell, but remained connected to the outer membrane (**Fig. 5h**). The graph in **Fig. 5i** shows the fluorescence intensity of the labeled (FAM-) DNA in **Fig. 5a–h**, during minutes 9-21 of the experiment (*cf.* **SI S5** for the graph showing the full time period and 19 compartments). The plots and their corresponding subcompartments are labeled with matching numbers in (**Fig. 5a–b** and **5i**). The plots in the blue spectrum indicate the subcompartments which take up DNA during superfusion and lose it rapidly when the flow is terminated. The subcompartments represented by the plots in the red spectrum encapsulate the DNA and maintain it, leading to the gradual increase in amount of DNA in the internal volume. The ability to maintain the internalized constituents is associated with the sealing of the initial membrane invaginations^18^. The gray regions in the background show the fluorescence intensity of the stock DNA suspension delivered by the microfluidic device (circle in dashed white line in **Fig. 5a**), over the duration of superfusion. Some DNA containing subcompartments disappear, or are relocated over time, which made further monitoring not possible. These instances are marked with an ‘x’ in **Fig. 5i**.

Transient pore- and defect-mediated encapsulation of external compounds in surface-adhered protocells has been reported both for small fluorescent dyes and genetic polymers^14, 18, 30^. In our previous work, we showed that subcompartments can also exchange compounds through the nano-sized openings connected to the proximal membrane^18^. Both uptake mechanisms are possibly involved in the DNA encapsulation shown in **Fig. 5**: On the one hand, we observed an instant loss of internal compounds when superfusion is terminated (plots in blue spectrum in **Fig. 5i**), on the other hand, we measured a gradual decrease in fluorescent intensity (plots in red spectrum in **Fig. 5i, SI Movie S3**).

**Fig. 5j–n** shows the dynamic exo-compartments which spontaneously emerge (**Fig. 5k, m**), uptake ambient DNA, and merge with the protocell superstructure (**Fig. 5l, n**) (*cf.* **SI Movie S3**). The graph in **Fig. 5o** shows the fluorescence intensity of different exo-compartments inside the superstructure, marked with an arrow in **Fig. 5j,** as well as the average intensity of the entire superstructure (black plot) over time. Colored plots represent different exo-compartments. The area under each plot is filled with solid color to emphasize the features of the plots. Some of the exo-compartments spontaneously disappear while no significant increase of DNA is observed inside the protocell (green t=0-10 min). After the merging of the other exo-compartments, a visible increase in DNA intensity inside the protocell was observed (asterisks in **Fig. 5o**). The gray zones show the time periods during which the DNA was externally exposed to the superstructures.

The extracellular vesicles, sometimes associated with nanotubes, are ubiquitous throughout all three domains of life^48, 49^. In eukaryotic cells, extracellular compartments are employed for cellular communication, e.g. during cell differentiation, mostly by formation of microvesicles with a size bracket up to 1 μm^50^. Formation of larger compartments from the plasma membrane, comparable in size with giant vesicles, have been reported in breast cancer cells^51^. In almost all cases, the extracellular vesicles are separated from the plasma membrane and released into the extracellular space. To our knowledge, the formation of membrane compartments that remain anchored the membrane has not been previously reported.

### DNA strand-displacement reactions

The multicompartmentalized protocell superstructures can encapsulate both single- and double-stranded DNA, and are able to accommodate strand displacement reactions (**Fig. 6**)^52^. We have selected this non-enzymatic process as an example for reactions involving prebiotically relevant molecules. It requires few reactants, and monitoring its outcome by confocal microscopy is straight-forward. Initially, the superstructures were grown in an aqueous environment containing dsDNA (**Fig. 6a**). The dsDNA consists of a fluorescently labeled strand which is initially quenched by its complementary strand, ‘the quencher’ featuring an overhanging region, i.e., a toehold (**Fig. 6b**). Subsequently, an invading strand which is complementary to the quencher strand of the dsDNA, was locally superfused with the superstructures. When the invading strand enters the protocell, it hybridizes with the quencher strand, causing melting of the initial dsDNA, followed by release of the fluorescently labeled strand and fluorescence emission (**Fig. 6b**). **Fig. 6c–e**, **Fig. 6f–h** shows the confocal micrograph time series of two different experiments corresponding to **Fig 6a–b**.

**Figure 6.**
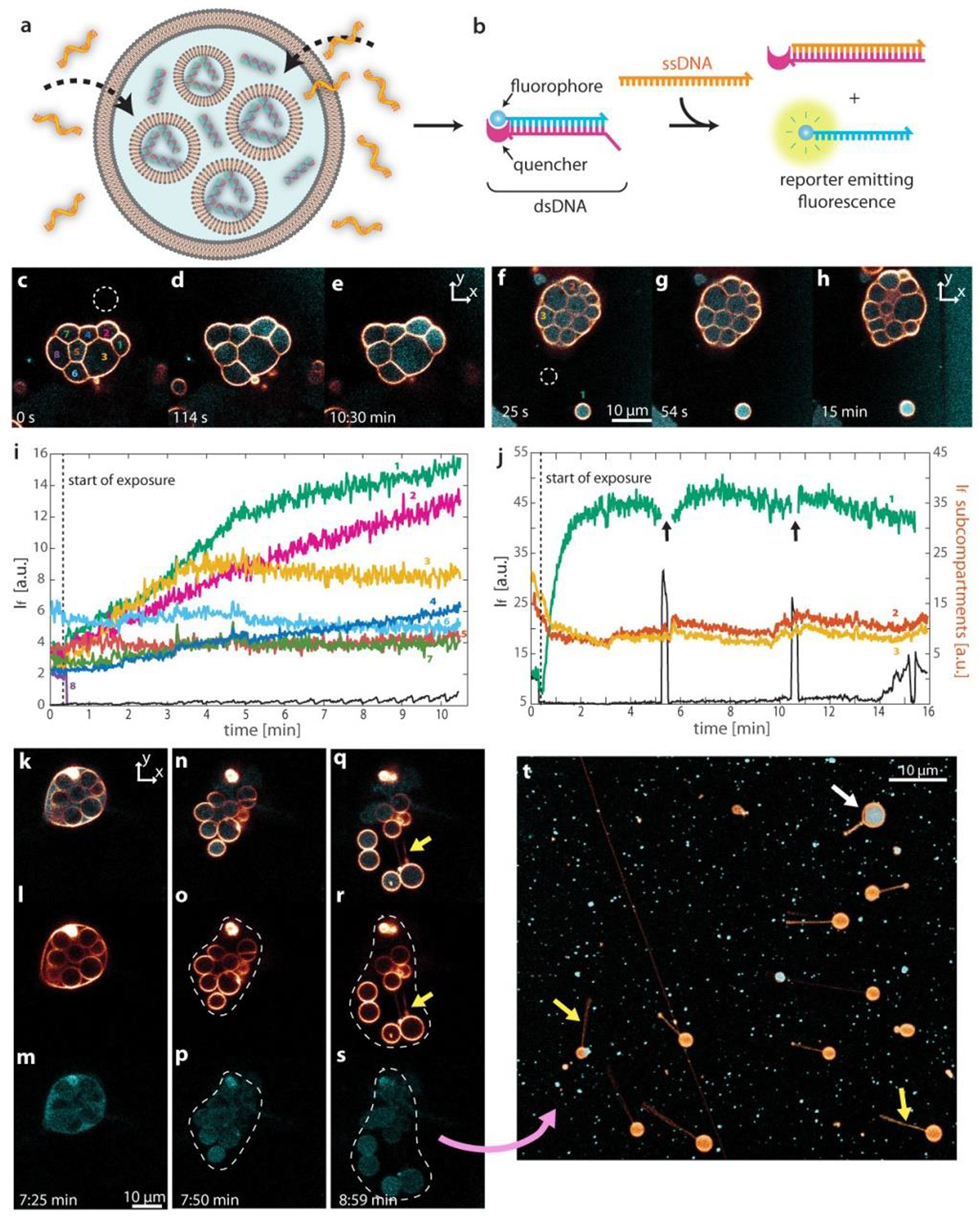
DNA strand displacement reactions and pseudo-division. (**a-b**) Schematic drawing of the displacement reaction. (**a**) A protocellular superstructure encapsulating dsDNA. The structure is exposed to ssDNA leading to its uptake. (**b**) ssDNA reacts with (the toe-hold of) the dsDNA inside the superstructures and hybridizes with the quencher DNA strand. The initially quenched fluorescent DNA strand will be released, leading to an increase in the measured fluorescence intensity. (**c-e**)-(**f-h**) Confocal micrograph time series showing two different experiments corresponding to (**a-b**). In (**f-h**) an isolated lipid compartment (marked as #1) is positioned next to the superstructure. (**i**) and (**j**) show fluorescence intensity vs. time, corresponding to (**c-e**) and (**f-h**). For each experiment, the fluorescence intensity of the ambient solution (black plots in i and j) was measured at the region in c and f encircled in dashed white lines. The numbers in the micrographs denominate regions of interest, the corresponding plots have matching numbers. (**k-t**) Pseudo-division of a superstructure encapsulating the reporter DNA. (**k-n-q**) DNA and membrane fluorescence channels as overlay. (**l-o-r**) Membrane fluorescence channel. (**m-p-s**) DNA fluorescence channel. (**t**) Daughter cells migrated and anchored to the surface. Yellow arrows point to the nanotubes connected to each daughter cell. The white arrow points to the daughter cell which has maintained its DNA cargo.

**Fig. 6i** shows the fluorescence intensity over time of each individual subcompartment inside the superstructure shown **Fig. 6c–e**, as well as a region outside the structure. The region of interest outside the protocell colony in which the intensity is monitored is encircled in white dashed lines in **Fig. 6c**, and its fluorescence intensity is represented by the black plot in **Fig. 6i**. The colored plots represent the intensity of the internal subcompartments. A subcompartment and its corresponding plot are denominated with matching numbers. Compartments 1 and 2 depict a gradually increasing fluorescence signal over the course of ~10 min.

**Fig. 6f–h** shows an example in which an isolated unilamellar vesicle (#1) is located next to the superstructure. In this case, the unilamellar vesicle appears to rapidly uptake the exchange strand over time, leading to a significantly increased fluorescence signal (green plot) while the protocell colony does not exhibit increasing fluorescence. This supports the earlier observations that the enveloping bilayer of the superstructures serves as an extra permeation barrier (**Fig. 4**). Note that the superstructure exhibits a low intensity fluorescence signal in its internal volume before the ssDNA was introduced. This could be due to the encapsulation of DNA which was not fully hybridized during annealing prior to the experiment. Over time the fluorescence intensity slightly increases during exposure to the ssDNA. The two time points at which the recirculation of ssDNA was stopped, can be observed from the black line plot in **Fig. 6j** (two spikes of increased fluorescence). When the recirculation of ssDNA is active, a hydrodynamically confined zone forms around the superstructure, free of any ambient fluorescent DNA. When the recirculation is terminated, the zone disappears leading to the increase in the fluorescence signal. These pulses can be observed in **SI Movie S4**. During pulsing, the single unilamellar vesicle in **Fig. 6f–h** occasionally moves out of the region of interest. At these moments the fluorescence intensity could not be properly collected (gaps in the green colored plot). Note further that the ssDNA exposure in **Fig. 6i** is continuous and in **Fig. 6j** has been applied in cycles.

### Pseudo-division of superstructures

**Fig. 6k–s** shows one superstructure undergoing pseudo-division^18^. This occurs when the enveloping membrane ruptures and the subcompartments migrate and eventually become free surface anchored daughter cells. **Fig. 6k–m** show the protocell structure 7 min 25 s after the DNA strand displacement experiment was initiated. **Fig. 6k–n–q** shows the lipid membrane and DNA fluorescence channels overlaid, **Fig. 6l–o–r** contains graphs of the membrane fluorescence only, and the graphs in **Fig. 6m–p–s** represent the DNA fluorescence only. A video of the original protocell bursting and releasing the daughter cells is provided as supplementary material (**SI Movie S4**). The daughter cells migrate and attach to other locations on the substrate. In **Fig. 6t** daughter cells anchored to the substrate with nanotethers are displayed. It is well-established that the vesicles exposed to a hydrodynamic force move and grow nanotethers^53^. Accordingly, the nanotubes we observe are most likely formed spontaneously during migration of daughter cells. Another possibility is that they maintain the nanotethers from the original intra-protocellular formation event (arrows in **Fig. 6q, r**). One daughter cell still carries its fluorescence content acquired prior to the pseudo-division event (white arrow in **Fig. 6t**). Similar structures have been reported in migrating eukaryotic cells, which leave long tubular strands terminated with large cargo vesicles (migrasomes)^54^. These connecting tubes eventually break, and migrasomes can transport the cytosolic content to other cells.

Giant vesicle compartments can act as miniature reactors and host various (bio)chemical systems, including intravesicular DNA strand melting and annealing^55^, reverse transcription PCR^56^, and light-triggered enzymatic reactions^57^. **Fig. 6** illustrates how the superstructures can encapsulate macromolecular DNA strands from the ambient environment and host a DNA displacement reaction^58^. This reaction was chosen as a relatively simple example of an enzyme-free prebiotic reaction similar to non-enzymatic RNA replication-related reactions^59, 60^.

## Conclusion

A tendency of biological cells towards aggregation is observed across all domains of life from bacterial colonies to multicellular organisms. The colony structure provides advantages, including greater physical stability and enhanced communication. Protocell colonies have been hypothesized as a possible development step towards the first cells^4, 10^.

We show a consistent pathway leading to colony-like protocell superstructures under simple conditions relevant to the early Earth. The presented system requires minimal set of initial components, such as amphiphilic molecules, a high energy surface, and a suitably composed aqueous environment. Subcompartments grow from the bottom-up inside a model protocell and exhibit high mechanical and osmotic stability. The protocell colony structures we produced in the laboratory can encapsulate genetic polymers from the ambient environment and host non-enzymatic reactions. Upon disintegration of the enveloping membrane, subcompartments detach and migrate, carrying the encapsulated cargo to remote locations. Membranous protocell superstructures which are able to uptake chemical compounds and maintain them in secluded spaces offer new possible pathways of development of prebiotic containers with an increasing set of functions and features towards primitive forms of life.

## Materials and Methods

### Preparation of lipid vesicles

The dehydration and rehydration method^61, 62^ was used to prepare the lipid suspensions. Briefly, lipids (99 wt %) and lipid-conjugated fluorophores (1 wt %) were dissolved in chloroform to a final concentration of 10 mg/ml (*cf.* **S1 Table** for a detailed list of lipid compositions). 300 μl of this mixture was then transferred to a 10 ml round bottom flask and the solvent was removed in a rotary evaporator at reduced pressure (20 kPa) for 6 hours to form a dry lipid film. The film was rehydrated with 3 ml of PBS buffer (5 mM Trizma Base, 30 mM K_3_PO_4_, 3 mM MgSO_4_.7H_2_O, 0.5 mM Na_2_EDTA, pH 7.4 adjusted with 1 M H_3_PO_4_) and stored at +4 °C overnight to allow the lipid cake to swell. The sample was then sonicated for 25 s at room temperature, leading to the formation of multi- and unilamellar giant vesicular compartments. For sample preparation, 4 μl of the lipid suspension was desiccated for 20 min, and the dry residue was subsequently rehydrated with 0.5 ml of HEPES-Na buffer containing 10 mM HEPES buffer and 100 mM NaCl, adjusted to pH 7.8 with 5 M NaOH. The lipid suspension was thereafter transferred onto a solid surface submerged in HEPES-Na buffer with addition of 4 mM CaCl2 (pH 7.8 adjusted with 5 M NaOH).

### Surface preparation

Aluminum (**Fig. 1–2, 4–6, Fig. S1**) surfaces were fabricated at the Microtechnology and Nanoscience facility at Chalmers University (MC2), Sweden or at the Norwegian Micro- and Nano-Fabrication Facility at the University of Oslo (MiNaLab). Al_2_O_3_ (**Fig. S1**) was deposited in MiNaLab. All thin films were deposited on glass cover slips (Menzel Gläss #1, 100-150 μm thickness; WillCo Wells B.V., Amsterdam, NL). Al films prepared at MC2 were deposited by sputter deposition using an FHR MS150 instrument (FHR Anlagenbau GmbH, Germany) to final thickness 10 nm. The substrates were pre-cleaned using ultrasonic bath of acetone for 10 minutes, followed by IPA and DI water bath, and blown dry with N2. Subsequently they were exposed to oxygen plasma cleaning (50 W, 250 mT, 2 min). At MiNa Labs, Al was deposited onto the glass substrates by E-beam evaporation using an EvoVac instrument (Ångstrom Engineering, Canada), also to a final thickness of 10 nm. Prior to the deposition the glass slides were cleaned using IPA, deionized water and blown dry with N_2_. Al_2_O_3_ was deposited onto glass substrates by atomic layer deposition (Beneq, Finland), to a final thickness of 10 nm. Surfaces were used immediately after their fabrication.

### Addition of chelators

To initiate the membrane instabilities leading to subcompartment formation, the buffer in the sample was gently exchanged with chelation buffer containing 10 mM HEPES, 100 mM NaCl, 10 mM EDTA and 7 mM BAPTA (pH 7.8, adjusted with 5 mM NaOH), using an automatic pipette 20 min after the initial deposition of the vesicles onto the substrates.

### Local heating

For the temperature control experiments shown in **Fig. 2**, an optical fiber coupled to IR-B laser was assembled to locally increase the temperature in the sample, in order to facilitate the de-wetting and subcompartment formation. A semiconductor diode laser (HHF-1470-6-95, λ = 1470 nm, Seminex), driven with an 8 A power source (4308 Laser Source, Arroyo Instruments) was used in combination with an 0.22 NA multimode optical fiber with 50 μm core diameter (Ocean Optics). The fiber exit was positioned approx. 30-50 μm from the vesicle by means of a micromanipulator. The laser current utilized for experiments were in the range 0.7-0-9 A, resulting in a local temperature increase to 40-70 °C^30^.

### Microfluidic device

The superfusion experiments shown in **Fig. 4–6** were performed using an open space microfluidic pipette (Fluicell AB, Sweden)^40^, positioned using a 3-axis water hydraulic micromanipulator (Narishige, Japan) in the vicinity of the protocell structures (30-50 μm distance). The structures were exposed to various solutes in pulses or in continuous flow, as specified for individual experiments. In the experiment illustrated in **Fig. S3** 25 μM fluorescein sodium salt in HEPES-Na buffer was used.

### Mechanical stability determination

In **Fig. 4** the model protocells and protocell superstructures were exposed to solutions of different osmotic strength (deionized water, HEPES-Na buffer) in intervals of 30 sec using the microfluidic device described above.

### Genetic polymers and DNA displacement reaction

The experiments involving DNA delivery shown in **Fig. 5** were performed using the microfluidic pipette described. The protocell superstructures were superfused with HEPES-Na buffer containing 25 μM of FAM-conjugated 20 base long DNA oligonucleotide (5’-/56-FAM/ TGT ACG TCA CAA CTA CCC CC-3’) (Dharmacon, USA) (pH 7.8, adjusted with 5 mM NaOH).

In the experiments shown in **Fig. 6** three DNA oligonucleotides, purified with HPLC (Integrated DNA technologies, USA), were used for the DNA displacement reaction reported by Zhang et al.^52^. The reactant sequences are as the following:

- reporter-ROX: /56-ROXN/CT TTC CTA CAC CTA CG
- reporter-RQ: TGG AGA CGT AGG TGT AGG AAA G/3IAbRQSp/
- output: CTT TCC TAC ACCTAC GTC TCC AAC TAA CTT ACG G

A thermal cycle was run to anneal 20 μM reporter-ROX and 20 μM reporter-RQ in TE buffer containing 10 mM Tris-HCl and 1mM EDTA (pH 8.0, adjusted with 5 mM NaOH and 1 mM HCl). The sample was incubated at 95 °C for 5 min and subsequently cooled to 20 °C over 75 min. The samples were further cooled and stored at +4 °C until the encapsulation experiment. To encapsulate DNA dimers (ROX-RQ) in the protocells and superstructures, DNA dimers were added to the chelation buffer to a final concentration of 5 μM during buffer exchange. After formation of structures, the excess dimers were removed by a second buffer exchange step, using dimer-free chelation buffer administered by an automatic pipette. 10 μM output strand in HEPES-Na buffer was locally delivered to the vicinity of structures in pulses, using the microfluidic pipette. The product of the DNA displacement reaction was observed as fluorescence signal produced by the released reporter-ROX strand.

### Microscopy imaging

All microscopy images were acquired with a laser scanning confocal microscopy system (Leica SP8, Germany) using a HCX PL APO CS 40x oil, NA 1.3 objective. The excitation/emission wavelengths varied with the employed fluorophores: Rhodamine B ex: 560 nm/em: 583 nm (**Fig. 1–2, 4–5**); ATTO 488 ex: 488 nm/em: 505 nm (**Fig. 6**), Texas Red DHPE ex: 595 nm/em: 615 (**Fig. S1)**; FAM-DNA ex: 488 nm, em: 515 nm (**Fig. 5**), ssDNA ex 588 nm/ em: 608 nm (**Fig. 6**).

### Image analyses

3D fluorescence micrographs were reconstructed using the Leica Application Suite X Software (Leica Microsystems, Germany). Image enhancement of fluorescence micrographs for the figures were performed with Adobe Photoshop CS4 (Adobe Systems, USA). The image analysis shown in **Fig. 1l,m** and fluorescence intensity analyses shown in **Fig. 5i,o**, **Fig. 6I,n**, were performed with the NIH Image-J software. Roundness as shown in **Fig. 1l,m** is defined as 4×area/(π×major_axis^2^) and roundness data for each histogram were normalized to a total number of subcompartments in each graph, and expressed in percentage. All graphs were plotted in Matlab R2018a. Schematic drawings were created with Adobe Illustrator CS4 (Adobe Systems, USA).

### Elastohydrodynamic theory for the membrane dynamics

We assume that at early time stages when the membrane deformations are small, the membrane height is given by *h*(*x*, *t*) = *h*_0_ + *h*_1_(*x*, *t*) with *h*_1_ ≪ *h*_0_ where *h*_0_ is the height of undeformed membrane. In the limit 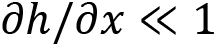, to a linear order in *h*_1_, the pressure difference *p* across the lipid membrane is given by,

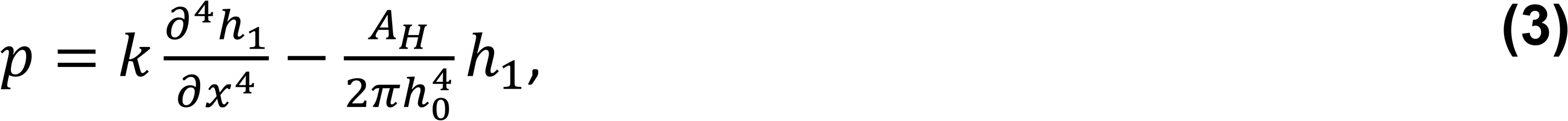

where *k*=10^−19^ J is the bending modulus of the membrane and *A*_*H*_=10^−21^ J is the Hamaker constant that characterizes the Van der Waals interactions between membrane and the aluminum substrate. The local mass conservation of the incompressible lipid membrane and the incompressible fluid underneath require that the rate of change of height *h* be governed by spatial variation of the liquid flux *hU* in the horizontal direction, where *U* is horizontal flow speed. To a linear order in *h*_1_, the mass conservation is given by of a thin liquid film

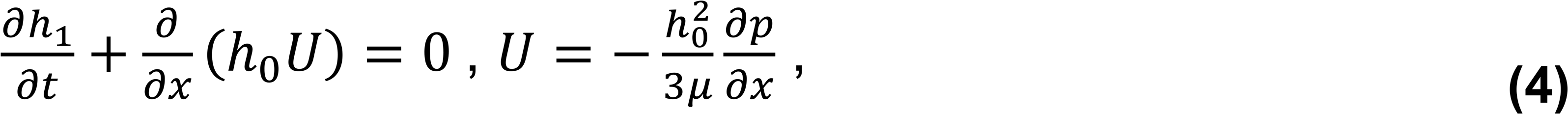

where μ is the dynamic viscosity of the solvent. **Eq. 4** along with **Eq. 3** constitute our elastohydrodynamic theory. Since the coefficients in the **Eq. 4** are independent of *x* and *t*, we explored the solutions of the form *h*_1_ = |*h*_1_|*e*^(*iqx*+*st*)^, where |*h*_1_| is the fluctuation amplitude at *t*=0 resulting from the thermal energy, *q* is the wave number of a given mode, and *s* is the corresponding inverse deformation time scale. In dimensionless units, substituting this relation into **Eq. 4**, we arrive at the following dispersion relation

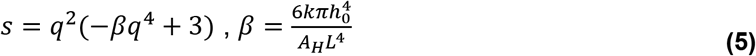

with critical wave number 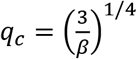. For numerical implementation we used the open source finite element analysis (FEA) library FEniCS on Python 3.6^63^. At *t*=0, the membrane height profile is given by *h*(*x*)=1+0.001cos(*qx*), and the time evolution is subject to the following boundary conditions: at *x*=0 we assume 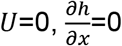 and 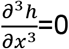; at *x*=1 we assume *h*=1, *p*=0 and 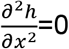.

## Acknowledgements

This work was made possible through financial support obtained from the Research Council of Norway (Forskningsrådet), Project Grant 274433, UiO: Life Sciences Convergence Environment, as well as the startup funding provided by the Centre for Molecular Medicine Norway (RCN 187615), and the Faculty of Mathematics and Natural Sciences at the University of Oslo. C.N.K. acknowledges the financial support by the College of Science at Virginia Tech. C. K. was partially supported by the Graduate School Doctoral Assistantship at Virginia Tech. R.R. gratefully acknowledges the H2020 ITN “Chemical Reaction Networks – CReaNET” – Ref. 812868. E.P.V. and R.R. thank the microtechnology center MC2 at Chalmers University of Technology for technical support.

## Notes

### Competing Interest Statement

The authors have declared no competing interest.

## References

1. Ganti, T., The Principles of Life. Oxford University Press UK: 2003.

2. Lopez, A.; Fiore, M., Investigating Prebiotic Protocells for A Comprehensive Understanding of the Origins of Life: A Prebiotic Systems Chemistry Perspective. Life (Basel, Switzerland) 2019, 9 (2), 49.

3. Lyu, Y.; Peng, R.; Liu, H.; Kuai, H.; Mo, L.; Han, D.; Li, J.; Tan, W., Protocells programmed through artificial reaction networks. Chemical Science 2020, 11 (3), 631–642.

4. Monnard, P.-A.; Walde, P., Current Ideas about Prebiological Compartmentalization. Life (Basel, Switzerland) 2015, 5 (2), 1239–1263.

5. Thompson, D. A. W., On growth and form. New ed. ed.; University Press ;: Cambridge :, 1945.

6. Carrara, P.; Stano, P.; Luisi, P. L., Giant Vesicles “Colonies”: A Model for Primitive Cell Communities. 2012, 13 (10), 1497–1502.

7. Casas-Ferrer, L.; Brisson, A.; Massiera, G.; Casanellas, L., Design of vesicle prototissues as a model for cellular tissues. Soft Matter 2021, 17 (19), 5061–5072.

8. Jin, L.; Kamat, N. P.; Jena, S.; Szostak, J. W., Fatty Acid/Phospholipid Blended Membranes: A Potential Intermediate State in Protocellular Evolution. Small 2018, 14 (15), 1704077.

9. de Souza, T. P.; Bossa, G. V.; Stano, P.; Steiniger, F.; May, S.; Luisi, P. L.; Fahr, A., Vesicle aggregates as a model for primitive cellular assemblies. Physical Chemistry Chemical Physics 2017, 19 (30), 20082–20092.

10. Wang, X.; Tian, L.; Du, H.; Li, M.; Mu, W.; Drinkwater, B. W.; Han, X.; Mann, S., Chemical communication in spatially organized protocell colonies and protocell/living cell micro-arrays. Chemical Science 2019, 10 (41), 9446–9453.

11. Bolognesi, G.; Friddin, M. S.; Salehi-Reyhani, A.; Barlow, N. E.; Brooks, N. J.; Ces, O.; Elani, Y., Sculpting and fusing biomimetic vesicle networks using optical tweezers. Nature Communications 2018, 9 (1).

12. Li, Q.; Li, S.; Zhang, X.; Xu, W.; Han, X., Programmed magnetic manipulation of vesicles into spatially coded prototissue architectures arrays. Nature Communications 2020, 11 (1), 232.

13. Bernard, A.-L.; Guedeau-Boudeville, M.-A.; Jullien, L.; di Meglio, J.-M., Raspberry vesicles. Biochimica et Biophysica Acta (BBA) - Biomembranes 2002, 1567, 1–5.

14. Köksal, E. S.; Põldsalu, I.; Friis, H.; Mojzsis, S.; Bizzarro, M.; Gözen, I., Spontaneous formation of prebiotic compartment colonies on Hadean Earth and pre-Noachian Mars. bioRxiv 2021, 2021.05.11.443509.

15. Akashi, K.; Miyata, H.; Itoh, H.; Kinosita, K. Jr., Formation of giant liposomes promoted by divalent cations: critical role of electrostatic repulsion. Biophysical journal 1998, 74 (6), 2973–2982.

16. Melcrová, A.; Pokorna, S.; Pullanchery, S.; Kohagen, M.; Jurkiewicz, P.; Hof, M.; Jungwirth, P.; Cremer, P. S.; Cwiklik, L., The complex nature of calcium cation interactions with phospholipid bilayers. Scientific Reports 2016, 6 (1), 38035.

17. Bilal, T.; Gözen, I., Formation and dynamics of endoplasmic reticulum-like lipid nanotube networks. Biomaterials Science 2017, 5 (7), 1256–1264.

18. Spustova, K.; Köksal, E. S.; Ainla, A.; Gözen, I., Subcompartmentalization and Pseudo-Division of Model Protocells. Small 2021, 17 (2), 2005320.

19. Gözen, I.; Dommersnes, P.; Czolkos, I.; Jesorka, A.; Lobovkina, T.; Orwar, O., Fractal avalanche ruptures in biological membranes. Nature Materials 2010, 9 (11), 908–912.

20. Kindt, J. T.; Szostak, J. W.; Wang, A., Bulk Self-Assembly of Giant, Unilamellar Vesicles. ACS Nano 2020, 14 (11), 14627–14634.

21. Zupanc, J.; Drašler, B.; Boljte, S.; Kralj-Iglič, V.; Iglič, A.; Erdogmus, D.; Drobne, D., Lipid vesicle shape analysis from populations using light video microscopy and computer vision. PloS one 2014, 9 (11), e113405–e113405.

22. Nadell, C. D.; Drescher, K.; Foster, K. R., Spatial structure, cooperation and competition in biofilms. Nature Reviews Microbiology 2016, 14 (9), 589–600.

23. Rao, M.; Eichberg, J.; Oró, J., Synthesis of phosphatidylcholine under possible primitive Earth conditions. Journal of Molecular Evolution 1982, 18 (3), 196–202.

24. Rao, M.; Eichberg, J.; Oró, J., Synthesis of phosphatidylethanolamine under possible primitive earth conditions. Journal of Molecular Evolution 1987, 25 (1), 1–6.

25. Hargreaves, W. R.; Mulvihill, S. J.; Deamer, D. W., Synthesis of phospholipids and membranes in prebiotic conditions. Nature 1977, 266 (5597), 78–80.

26. Liu, L.; Zou, Y.; Bhattacharya, A.; Zhang, D.; Lang, S. Q.; Houk, K. N.; Devaraj, N. K., Enzyme-free synthesis of natural phospholipids in water. Nature Chemistry 2020, 12 (11), 1029–1034.

27. Toner, J. D.; Catling, D. C., A carbonate-rich lake solution to the phosphate problem of the origin of life. Proceedings of the National Academy of Sciences of the United States of America 2020, 117 (2), 883–888.

28. Kazmierczak, J.; Kempe, S.; Kremer, B., Calcium in the early evolution of living systems: A biohistorical approach. Current Organic Chemistry 2013, 17 (16), 1738–1750.

29. Hashizume, H., Role of Clay Minerals in Chemical Evolution and the Origins of Life. In Clay Minerals in Nature, Intech Open: 2012.

30. Köksal, E. S.; Liese, S.; Xue, L.; Ryskulov, R.; Viitala, L.; Carlson, A.; Gözen, I., Rapid growth and fusion of protocells in surface-adhered membrane networks. 2020, 2020.03.10.980417.

31. Feng, J.; He, Y., Collective motion of bacteria and their dynamic assembly behavior. SCIENCE CHINA Materials 2017, 60 (2095-8226), 1079.

32. Zhang, H. P.; Be’er, A.; Florin, E. L.; Swinney, H. L., Collective motion and density fluctuations in bacterial colonies. Proceedings of the National Academy of Sciences 2010, 107 (31), 13626–13630.

33. Langevin, D., Aqueous foams and foam films stabilised by surfactants. Gravity-free studies. Comptes Rendus Mécanique 2017, 345 (1), 47–55.

34. Winkelmann, J.; Dunne, F. F.; Langlois, V. J.; Möbius, M. E.; Weaire, D.; Hutzler, S., 2D foams above the jamming transition: Deformation matters. Colloids and Surfaces A: Physicochemical and Engineering Aspects 2017, 534, 52–57.

35. Steinkühler, J.; Knorr, R. L.; Zhao, Z.; Bhatia, T.; Bartelt, S. M.; Wegner, S.; Dimova, R.; Lipowsky, R., Controlled division of cell-sized vesicles by low densities of membrane-bound proteins. Nature Communications 2020, 11 (1), 905.

36. Phillips, R.; Kondev, J.; Theriot, J.; Garcia, H. G.; Orme, N., Physical biology of the cell. 2013.

37. Israelachvili, J. N., 6 - Van der Waals Forces. In Intermolecular and Surface Forces (Third Edition), Israelachvili, J. N., Ed. Academic Press: San Diego, 2011; pp 107–132.

38. Oron, A.; Davis, S. H.; Bankoff, S. G., Long-scale evolution of thin liquid films. Reviews of Modern Physics 1997, 69 (3), 931–980.

39. Chaurasia, A. K.; Rukangu, A. M.; Philen, M. K.; Seidel, G. D.; Freeman, E. C., Evaluation of bending modulus of lipid bilayers using undulation and orientation analysis. Physical Review E 2018, 97 (3), 032421.

40. Ainla, A.; Jeffries, G. D. M.; Brune, R.; Orwar, O.; Jesorka, A., A multifunctional pipette. Lab on a Chip 2012, 12 (7), 1255–1261.

41. Ho, J. C. S.; Rangamani, P.; Liedberg, B.; Parikh, A. N., Mixing Water, Transducing Energy, and Shaping Membranes: Autonomously Self-Regulating Giant Vesicles. Langmuir 2016, 32 (9), 2151–2163.

42. Gözen, İ., Did Solid Surfaces Enable the Origin of Life? Life 2021, 11 (8).

43. Bhatia, T.; Agudo-Canalejo, J.; Dimova, R.; Lipowsky, R., Membrane Nanotubes Increase the Robustness of Giant Vesicles. ACS Nano 2018, 12 (5), 4478–4485.

44. Vanhille-Campos, C.; Šarić, A., Modelling the dynamics of vesicle reshaping and scission under osmotic shocks. Soft Matter 2021, 17 (14), 3798–3806.

45. Schrum, J. P.; Zhu, T. F.; Szostak, J. W., The origins of cellular life. Cold Spring Harbor perspectives in biology 2010, 2 (9), a002212–a002212.

46. Gilbert, W., Origin of life: The RNA world. Nature 1986, 319 (6055), 618–618.

47. Saha, R.; Pohorille, A.; Chen, I. A., Molecular Crowding and Early Evolution. Origins of Life and Evolution of Biospheres 2014, 44 (4), 319–324.

48. Gill, S.; Catchpole, R.; Forterre, P., Extracellular membrane vesicles in the three domains of life and beyond. FEMS microbiology reviews 2019, 43 (3), 273–303.

49. Johansen, J.; Ramanathan, V.; Beh, C. T., Vesicle trafficking from a lipid perspective: Lipid regulation of exocytosis in Saccharomyces cerevisiae. Cellular logistics 2012, 2 (3), 151–160.

50. Ratajczak, M. Z.; Ratajczak, J., Extracellular microvesicles/exosomes: discovery, disbelief, acceptance, and the future? Leukemia 2020, 34 (12), 3126–3135.

51. Wright, P. K.; Jones, S. B.; Ardern, N.; Ward, R.; Clarke, R. B.; Sotgia, F.; Lisanti, M. P.; Landberg, G.; Lamb, R., 17β-estradiol regulates giant vesicle formation via estrogen receptor-alpha in human breast cancer cells. Oncotarget 2014, 5 (10).

52. Zhang, D. Y.; Turberfield, A. J.; Yurke, B.; Winfree, E., Engineering entropy-driven reactions and networks catalyzed by DNA. Science 2007, 318 (5853), 1121–5.

53. Borghi, N.; Kremer, S.; Askovic, V.; Brochard-Wyart, F., Tube extrusion from permeabilized giant vesicles. Europhysics Letters (EPL) 2006, 75 (4), 666–672.

54. Ma, L.; Li, Y.; Peng, J.; Wu, D.; Zhao, X.; Cui, Y.; Chen, L.; Yan, X.; Du, Y.; Yu, L., Discovery of the migrasome, an organelle mediating release of cytoplasmic contents during cell migration. Cell Research 2015, 25 (1), 24–38.

55. Mansy, S. S.; Szostak, J. W., Thermostability of model protocell membranes. Proceedings of the National Academy of Sciences 2008, 105 (36), 13351–13355.

56. Tsugane, M.; Suzuki, H., Reverse Transcription Polymerase Chain Reaction in Giant Unilamellar Vesicles. Scientific Reports 2018, 8 (1), 9214.

57. Hindley, J. W.; Elani, Y.; McGilvery, C. M.; Ali, S.; Bevan, C. L.; Law, R. V.; Ces, O., Light-triggered enzymatic reactions in nested vesicle reactors. Nature Communications 2018, 9 (1), 1093.

58. Zhang, D. Y.; Winfree, E., Control of DNA Strand Displacement Kinetics Using Toehold Exchange. Journal of the American Chemical Society 2009, 131 (47), 17303–17314.

59. Kim, S. C.; Zhou, L.; Zhang, W.; O’Flaherty, D. K.; Rondo-Brovetto, V.; Szostak, J. W., A Model for the Emergence of RNA from a Prebiotically Plausible Mixture of Ribonucleotides, Arabinonucleotides, and 2′-Deoxynucleotides. Journal of the American Chemical Society 2020, 142 (5), 2317–2326.

60. Zhang, S. J.; Duzdevich, D.; Szostak, J. W., Potentially Prebiotic Activation Chemistry Compatible with Nonenzymatic RNA Copying. Journal of the American Chemical Society 2020, 142 (35), 14810–14813.

61. Köksal, E. S.; Belletati, P. F.; Reint, G.; Olsson, R.; Leitl, K. D.; Kantarci, I.; Gözen, I., Spontaneous Formation and Rearrangement of Artificial Lipid Nanotube Networks as a Bottom-Up Model for Endoplasmic Reticulum. JoVE 2019, (143), e58923.

62. Karlsson, M.; Nolkrantz, K.; Davidson, M. J.; Strömberg, A.; Ryttsén, F.; Åkerman, B.; Orwar, O., Electroinjection of Colloid Particles and Biopolymers into Single Unilamellar Liposomes and Cells for Bioanalytical Applications. Analytical Chemistry 2000, 72 (23), 5857–5862.

63. Alnæs, M. S.; Blechta, J.; Hake, J.; Johansson, A.; Kehlet, B.; Logg, A.; Richardson, C.; Ring, J.; Rognes, M. E.; Wells, G. N., The FEniCS project version 1.5. Archive of Numerical Software 2015, (Vol. 3, No. 100).

