## Supporting Information for "Colony-like Protocell Superstructures"

### S1. Lipid compositions and surfaces

**Table S1.** Lipid, lipid-conjugated fluorophore and solid surface combinations

<table>
<tr><th>Surface</th><th>Lipid species</th><th>wt %</th><th>Fluorophore (1%)</th><th>Associated figure</th></tr>
<tr><td rowspan="4">Al</td><td rowspan="2">E.coli Pol. Ext.</td><td rowspan="2">99</td><td>16:0 Liss Rhod B</td><td>Fig. 1-4, S2, S4</td></tr>
<tr><td>ATTO 488-DHPE</td><td>Fig. 5</td></tr>
<tr><td>PE:PG:CA</td><td>67:23:9</td><td>16:0 Liss Rhod B</td><td>Fig. S1a, S3</td></tr>
<tr><td>PC:DOPE</td><td>69:30</td><td>Texas Red-DHPE</td><td>Fig. S1b</td></tr>
<tr><td>$Al_2O_3$</td><td>E.coli Pol. Ext.</td><td>99</td><td>16:0 Liss Rhod B</td><td>Fig. S1c</td></tr>
</table>

Abbreviations used in the table are as following:

| | |
|---|---|
| E.coli Pol. Ext.: | E.coli polar lipid extract |
| PE: | L-α-phosphoethanolamine (E.coli) |
| PG: | L-α –phosphatidylglycerol (E.coli) |
| CA: | Cardiolipin (E.coli) |
| PC: | L-α-phosphatidylcholine (Soy) |
| DOPE: | 18:1 Dioleoylphosphoethanolamine |

All lipid products and 16:0 Liss Rhodamine PE were purchased from Avanti Polar Lipids, USA. ATTO 488-DHPE was purchased from ATTO-TEC Gmbh, Germany. Texas Red-DHPE was purchased from Sigma-Aldrich, USA.

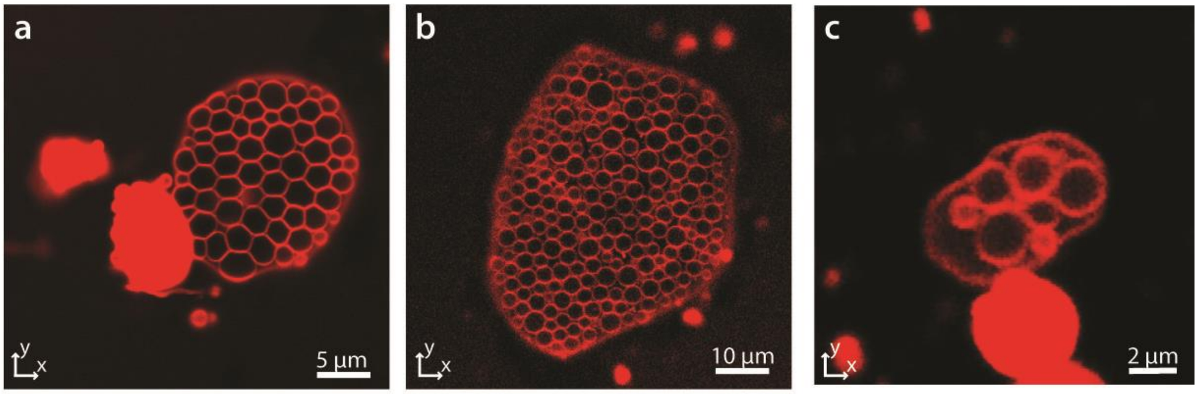

**Figure S1.** Protocell colonies made from (**a**) PE-PG-CA, and (**b**) PC-DOPE on Al surface, and from (**c**) E.coli Pol. Ext. on $Al_2O_3$ surface.

### S2. Pseudo-division

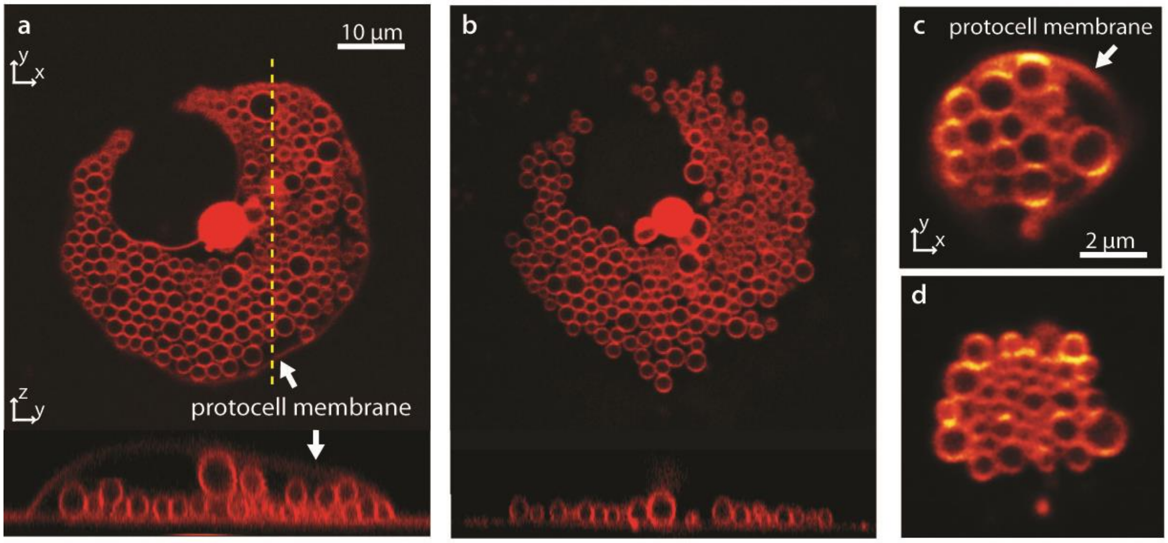

**Figure S2.** Pseudo-division of protocell superstructures. (**a-d**) Confocal micrographs of two superstructures before (**a, c**) and after pseudo-division event (**b, d**). The daughter protocells remain intact after the enveloping protocell membrane disintegrate. Superstructure membranes were made from E.coli Polar Extract lipids and positioned on Al surface.

### S3. Continuum theory for the onset of the subcompartment formation

#### S3.1 Elastohydrodynamic instability as a plausible mechanism for the birth of subcompartments

Removing divalent calcium ions that pin vesicles of negatively charged lipid bilayers on a negatively charged aluminum surface results in the formation of subcompartmental bulges at the membrane-substrate interface in an aqueous ionic solvent (**Fig. S3.1, Fig. 3a** main article). We hypothesized that the growth of these subcompartments must be due to an elastohydrodynamic instability that may arise from three types of interactions at the interface:

1. Spontaneous curvature, which may emerge due to the asymmetry of lipid molecules or local density differences between the monolayers that constitute the bilayer,

2. Electrostatic repulsions between the like-charged membrane and substrate, which must be weakened due to the ionic screening of the solvent,

3. Attractive van der Waals (vdW) interactions across the substrate-solvent-bilayer system that is at play at a $\lesssim 100$ nm range.

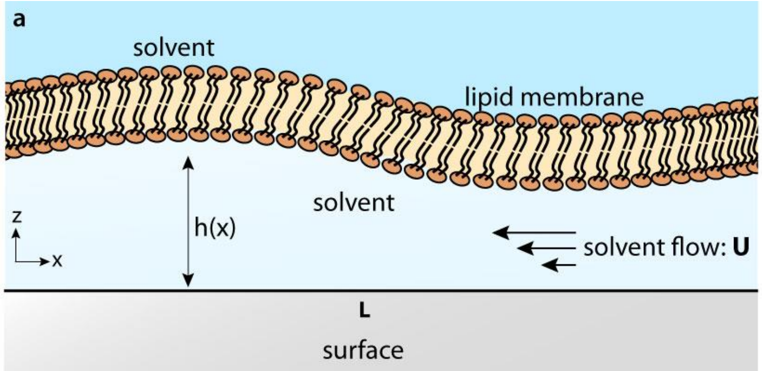

**Figure S3.1: The bilayer-solvent-substrate system.** At the ventral region of the membrane, the lipid bilayer and the aluminum substrate confine a thin film of aqueous ionic background solvent, which flows in response to the pressure gradients along the $x$ −direction. The center point at the bottom of the vesicle corresponds to the left boundary, and the right boundary is located in the proximity to the vesicle front.

In the absence of electrostatic and van der Waals interactions, the equilibrium conformation of the lipid bilayer is determined by a balance between the local membrane bending characterized by the bending modulus $k$ (units: Joules), membrane-solvent interfacial tension $\gamma$ (units: Joules/area), the membrane spontaneous curvature $c_0$ , and the liquid pressure difference $p$ across the lipid membrane. Defining the membrane mean curvature as $H$ and the membrane Gaussian curvature as $\kappa_G$ , the local stress balance on the membrane is then given as[1-2]

$$2k\nabla^2 H + k(2H + c_0)(2H^2 - 2\kappa_G - c_0 H) - 2\gamma H = p\,. \quad (1)$$

When the membrane is in the close proximity to a surface at a nanometer scale, the stress balance in the bulk, given by Eq. (1), must be modified to take into account the electrostatic interactions and vdW interactions between the membrane and the substrate. When the membrane conformation undergoes small vertical undulations, the membrane electrostatic potential energy density per unit area $dU_q$ will have a variation $\delta dU_q$ . Accordingly, we will label the associated stresses as $\delta du_q$ . When the Hamaker constant of the substrate-solvent-bilayer system $A_H$ is known, the vdW interactions due to the different polarizabilities of the lipid, solvent, and aluminum molecules can be modeled by noting that the ventral region of the vesicle approximately conforms with the flat substrate. Denoting the areal vdW interaction energy density by $dU_w$ and the associated stresses by $\delta du_w$ , Eq. (1) becomes in the presence of electrostatic and van der Waals contributions

$$2k\nabla^2 H + k(2H + c_0)(2H^2 - 2\kappa_G - c_0 H) - 2\gamma H + \delta du_q + \delta du_w = p\,. \quad (2)$$

For simplicity, we look at the two-dimensional (2D) cross section of the ventral region of the vesicle and assume that chelation happens uniformly along the horizontal direction (**Fig. S3.1**). Since the radius of the vesicle $L$ is much larger than the distance $h$ between the lipid membrane and the aluminum substrate, the slope of the bilayer profile must satisfy $\partial h/\partial x \ll 1$ at the onset of membrane detachment. In this limit, the mean curvature of the 2D membrane cross-section can be simplified to $H \approx \frac{1}{2}\partial^2 h/\partial x^2$ with a vanishing Gaussian curvature $\kappa_G = 0$. Note that because a lipid bilayer is in general subject to big thermal fluctuations yet does not undergo spontaneous division when $c_0 = 0$, it can be estimated that $\gamma \approx 0$ in Eq. (2). Then, Eq. (2) becomes

$$k\frac{\partial^4 h}{\partial x^4} + \frac{k}{2}\left(\frac{\partial^2 h}{\partial x^2}\right)^3 - \frac{c_0^2 k}{2}\frac{\partial^2 h}{\partial x^2} + \delta du_q + \delta du_w = p\,. \tag{3}$$

Here, because the ions dissolved in the background liquid will screen the electrostatic interactions between the membrane and the surface, the resulting electrostatic pressure $\delta dU_q$ is determined by the Debye-Hückel theory when the electrostatic potential energy is much smaller than the thermal energy, i.e. $U_q \ll k_B T$[3]. The strength of electrostatic screening is governed by the Debye screening length $\lambda_D$, which depends on the electric permittivity of the aqueous medium, thermal energy, as well as the valency and concentration of the ionic species. If $h \gg \lambda_D$ ($h$: membrane height), electrostatic stresses are negligible. In our system $\lambda_D \approx 0.84$ nm and $h \sim 10$ nm, therefore, we retain this term to assess its stability properties.

As soon as the membrane is detached from the substrate upon chelation, the thermal energy $k_B T$ ($k_B$: Boltzmann's constant, $T$: temperature) results in fluctuations in the ventral lipid membrane height $h$. The amplitude of these height undulations is determined by $k$, provided that the equipartition theorem holds. When the membrane-liquid surface tension vanishes ($\gamma \approx 0$), the equipartition theorem yields for a Fourier mode with a wave number $q$

$$\langle |h(q)|^2 \rangle = \frac{k_B T}{L^2 k q^4}\,, \tag{4}$$

where $k_B T \sim 4 \times 10^{-21}$ J at room temperature. Whether these undulations will be amplified or suppressed depends on the dynamics of electrostatic interactions, van der Waals interactions and/or the effect of spontaneous curvature. In the following, first, we calculate the stresses emerging from electrostatic interactions and vdW interactions. Then we introduce the elastohydrodynamic thin film theory for the dynamics of the bilayer-liquid-substrate domain. Finally, based on our model, we use linear stability analysis to investigate the role of each of the interactions 1−3 in the emergence of the elastohydrodynamic instability.

#### S3.2 Stresses due to the electrostatic interactions

For $h \ll L$, the negatively charged lipid bilayer membrane and negatively charged aluminum surface can be modeled as two nearly parallel surfaces with an aqueous ionic solvent in between (**Fig. S3.1**). The screened electric field $E$ applied by the metal surface on the bilayer in terms of the membrane height $z = h$ is given by the Debye-Hückel theory (when $U_q \ll k_B T$ ; see Section 6 for mathematical details) in terms of the Debye length $\lambda_D$ ($\lambda_D \sim 0.84$ nm for our system),

$$E(h) = \frac{\sigma_s}{D\epsilon_0} e^{-\frac{h}{\lambda_D}} + \frac{\sigma_m}{2D\epsilon_0}(e^{-\frac{2h}{\lambda_D}} - 1)\,, \tag{5}$$

where $\sigma_s$ is the charge density of the aluminum surface, $\sigma_m$ is the charge density of the lipid membrane, $\epsilon_0$ is the vacuum permittivity, and $D$ is the dielectric constant of the ionic solvent. The corresponding electrostatic potential $V(h)$ is (see **S3.6**)

$$\begin{aligned} V(h) \quad &= \frac{\lambda_D \sigma_s}{D\epsilon_0} e^{-\frac{h}{\lambda_D}} + \frac{\sigma_m \lambda_D}{2D\epsilon_0} e^{-\frac{2h}{\lambda_D}} + \frac{\sigma_m \lambda_D}{2D\epsilon_0} \\ &= E(h)\lambda_D + \frac{\sigma_m}{D\epsilon_0}\lambda_D\,. \end{aligned} \tag{6}$$

On a membrane area element $dA$, the electrostatic potential energy $dU_q$ is then given by

$$dU_q = V\sigma_m dA = (E\sigma_m \lambda_D + v_0 \sigma_m)dA\,, \quad v_0 = \frac{\sigma_m \lambda_D}{D\epsilon_0}\,. \tag{7}$$

Locally, if the membrane height changes infinitesimally by $\delta h \equiv \bar{\delta}$ where $\bar{\delta} \ll h$, the variation in the potential energy $\delta dU_q$ becomes

$$\delta dU_q = \delta E \sigma_m \lambda_D dA + (E\sigma_m \lambda_D + v_0 \sigma_m)\delta dA \tag{8}$$

with

$$\delta dA = -2H\bar{\delta} dA + \mathcal{O}(\bar{\delta}^2)\,, \tag{9}$$

where $H$ is the mean curvature. The variation in the electric field $\delta E$ is then given by

$$\delta E = \frac{\partial E}{\partial h}\delta h = -\frac{\sigma_s \bar{\delta}}{D\epsilon_0 \lambda_D} e^{-\frac{h}{\lambda_D}} - \frac{\sigma_m \bar{\delta}}{D\epsilon_0 \lambda_D} e^{-\frac{2h}{\lambda_D}}\,. \tag{10}$$

The variation in the potential energy is rewritten as

$$\begin{aligned} \delta dU_q \quad &= \left[-\frac{\sigma_m}{D\epsilon_0}(\sigma_s e^{-\frac{h}{\lambda_D}} + \sigma_m e^{-\frac{2h}{\lambda_D}}) - 2H(E\sigma_m \lambda_D + v_0 \sigma_m)\right]\bar{\delta} dA \\ &\equiv \delta du_q \bar{\delta} dA. \end{aligned} \tag{11}$$

Using $H \approx \frac{1}{2}\partial^2 h/\partial x^2$ , the electrostatic stress $\delta du_q$ is given by

$$\delta du_q = -\frac{\sigma_m}{D\epsilon_0}(\sigma_s e^{-\frac{h}{\lambda_D}} + \sigma_m e^{-\frac{2h}{\lambda_D}}) - \frac{\partial^2 h}{\partial x^2}(E\sigma_m \lambda_D + v_0 \sigma_m)\,. \tag{12}$$

We will investigate the bilayer stability in the presence of the electrostatic stresses $\delta du_q$ in Section **S3.4** and show that $\delta du_q$ stabilizes the bilayer profile while being largely suppressed by the screening charges dissolved in the ionic solution.

**S3.3 Stresses due to van der Waals interactions**

The van der Waals interaction energy per unit area between two parallel interfaces is given by[4-5],

$$w(h) = -\frac{A_H}{12\pi h^2}. \tag{13}$$

For our bilayer-solvent-substrate system the Hamaker constant is calculated from the Lifshitz theory as $A_H = 2.08 \times 10^{-21}\, J$ [4].The vdW energy is given by $dU_w = wdA$ , and its variation due to infinitesimal height changes $\bar{\delta}$ yields

$$\delta dU_w = \frac{A_H}{6\pi h^3}\bar{\delta}dA + \frac{A_H}{6\pi h^2}H\bar{\delta}dA \tag{14}$$

Therefore, the stress on the lipid membrane due to Van der Waals interactions can be found as

$$\delta du_w = \frac{A_H}{6\pi h^2}\left[\frac{1}{h} + H\right]. \tag{15}$$

We will investigate the bilayer stability in the presence of the vdW stresses $\delta du_w$ in Section **S3.4** and show that for attractive interactions ($A_H > 0$), the membrane height is unstable above a critical wavelength set by the balance between the bending energy scale $k$ and the vdW energy scale $A_H$ .

**S3.4 Elastohydrodynamic thin film theory**

To quantify the onset of the formation of subcompartmental bulges at the bottom of a lipid bilayer vesicle, we have developed an elastohydrodynamic thin film theory that relates the coupling between the membrane profile and the mechanisms (1)-(3) (see **S3.1**) to the pressure gradients, which drive fluid flow underneath the membrane in a high-aspect-ratio domain characterized by $h \ll L$ (**Fig. S3.1**). The flow properties of thin liquid films can be modeled by the lubrication approximation in the limit $h \ll L$ that leads to $\partial h/\partial x \ll 1$[6]. The local mass conservation of the incompressible lipid membrane and the incompressible liquid underneath require that the rate of change of the membrane height $h$ be governed by the spatial variation of the liquid flux $hU$ in the horizontal direction ($U$: horizontal flow speed; see **Fig. S3.1**, $\mu$: liquid dynamic viscosity; units: $Pa \cdot s$), that is

$$\frac{\partial h}{\partial t} + \frac{\partial}{\partial x}(hU) = 0\,, \quad U = -\frac{h^2}{3\mu}\frac{\partial p}{\partial x}. \tag{16}$$

The mass conservation (Eq. (16)) and the liquid pressure (Eq. (3)) constitute our elastohydrodynamic model. Starting from a sinusoidal height profile for a given wave number $q$ and a corresponding amplitude set by Eq. (4), the model can be analyzed by using linear stability analysis that yields the dynamics of $h$ at early times. Substituting $\delta du_q$ (Eq. (12)) and $\delta du_w$ (Eq. (15)) in Eq. (3), we can write a complete expression for the stress balance on the lipid bilayer membrane as

$$k\frac{\partial^4 h}{\partial x^4}+\frac{k}{2}\left(\frac{\partial^2 h}{\partial x^2}\right)^3 \quad -\frac{c_0^2 k}{2}\frac{\partial^2 h}{\partial x^2}-\frac{\sigma_m}{D\epsilon_0}\left(\sigma_s e^{-\frac{h}{\lambda_D}}+\sigma_m e^{-\frac{2h}{\lambda_D}}\right)$$
$$-\frac{\partial^2 h}{\partial x^2}\left(E\sigma_m\lambda_D+v_0\sigma_m\right)+\frac{A_H}{6\pi h^2}\left[\frac{1}{h}+H\right]=p\,. \quad (17)$$

To make Eq. (17) dimensionless, we introduce the height, length, and pressure scales of the system (primes denote the dimensionless variables) as

$h \equiv h_0 h'$ $\quad$ $h_0$: initial height of the membrane; $h_0 = 10$nm,

$x \equiv Lx'$ $\quad$ $L$: radius of the vesicle; $L = 1\mu$m,

$p \equiv p_0 p'$ $\quad$ $p_0$: pressure scale.

Substituting above relations into Eq. (17) and then dropping the primes we obtain the stress balance equation in terms of dimensionless variables

$$\frac{kh_0}{L^4}\frac{\partial^4 h}{\partial x^4}+\frac{1}{2}\frac{kh_0^3}{L^6}\left(\frac{\partial^2 h}{\partial x^2}\right)^3-\frac{c_0^2 k h_0}{2L^2}\frac{\partial^2 h}{\partial x^2}$$
$$-\frac{\sigma_m^2}{D\epsilon_0}\left(\frac{\sigma_s}{\sigma_m}e^{-\frac{h_0 h}{\lambda_D}}+e^{-\frac{2h_0 h}{\lambda_D}}\right)-\frac{\sigma_m^2}{D\epsilon_0}\frac{\lambda_D h_0}{L^2}\frac{\partial^2 h}{\partial x^2}\left(\tilde{E}+1\right)+\frac{A_H}{6\pi h_0^2 h^2}\left[\frac{1}{h_0 h}+\frac{h_0}{2L^2}\frac{\partial^2 h}{\partial x^2}\right]=p_0 p\,, \quad (18)$$

with

$$\tilde{E}\equiv\frac{ED\epsilon_0}{\sigma_m}=\frac{\sigma_s}{\sigma_m}e^{-\frac{h_0 h}{\lambda_D}}+\frac{1}{2}\left(e^{-\frac{2hh_0}{\lambda_D}}-1\right).$$

Now we define the pressure scale $p_0 \equiv \frac{A_H}{6\pi h_0^3}$, the domain aspect ratio $\epsilon \equiv \frac{h_0}{L} \sim 10^{-2}$ ($\epsilon \ll 1$), the reduced bending modulus $\beta \equiv \frac{6k\pi\epsilon^4}{A_h} \sim 10^{-5}$, the reduced squared spontaneous curvature $\alpha \equiv \frac{3\pi k c_0^2 \epsilon^2 h_0^2}{A_H}$ ($\alpha \geq 0$), $\eta \equiv \frac{h_0}{\lambda_D}$, $\bar{\sigma} = \frac{\sigma_s}{\sigma_m} \approx 0.1$ and the reduced electrostatic stress coefficient $\zeta \equiv \frac{6\pi\sigma_m^2 h_0^3}{D\epsilon_0 A_h} \sim 10^3$. Dividing both sides of Eq. (18) by $p_0$, substituting the above constants, and retaining the the terms up to the order $\mathcal{O}(\epsilon^2)$, we obtain the dimensionless stress balance relation at the leading order

$$\beta\frac{\partial^4 h}{\partial x^4}-\alpha\frac{\partial^2 h}{\partial x^2}+\frac{1}{h^3}-\zeta\left(\bar{\sigma}e^{-\eta h}+e^{-2\eta h}\right)=p\,. \quad (19)$$

Here, the first term on the left-hand side is due to the bending energy, the second term due to the spontaneous curvature, the third term due to the vdW interactions, and the fourth term models the electrostatic interactions.

To derive the dimensionless form of the Eq. (16), we set the characteristic velocity scale $U_0 \equiv \frac{h_0^2 p_0}{3\mu L} \sim 10^{-5} m/s$. This return the time scale of longitudinal flows as $\tau \equiv L/U_0 \sim 0.1s$. By substituting $h \equiv h_0 h'$, $x \equiv Lx'$, $p \equiv p_0 p'$ and $t \equiv \tau t'$ in Eq. (16) and dropping the primes from the dimensionless variables, the dimensionless form of the thin film continuity equation (Eq. (16)) is given as

$$\frac{\partial h}{\partial t} - \frac{\partial}{\partial x}\left(h^3 \frac{\partial p}{\partial x}\right) = 0\,. \quad (20)$$

Substituting Eq. (19) into Eq. (20), the time evolution equation for the bilayer height $h$ reads

$$\frac{\partial h}{\partial t} - \frac{\partial}{\partial x}\left\{h^3 \frac{\partial}{\partial x}\left[\beta \frac{\partial^4 h}{\partial x^4} - \alpha \frac{\partial^2 h}{\partial x^2} + \frac{1}{h^3} - \zeta(\bar{\sigma} e^{-\eta h} + e^{-2\eta h})\right]\right\} = 0\,. \quad (21)$$

The linear stability analysis for small undulations about a constant bilayer height $h_0$ can be performed by considering the plane wave deformations

$$h = h_0 + h_1 \equiv h_0 + |h_1| e^{(iqx+st)}\,, \quad |h_1| \ll h_0\,, \quad (22)$$

where $s$ (units: 1/time) is the characteristic deformation rate with a wave number $q$ . Then, to linear order in $h_1$ , the nonlinear terms in Eq. (21) can be simplified as $h^3 \frac{\partial^5 h}{\partial x^5} \approx h_0^3 \frac{\partial^5 h_1}{\partial x^5}$, $h^3 \frac{\partial^3 h}{\partial x^3} \approx h_0^3 \frac{\partial^3 h_1}{\partial x^3}$, $\frac{1}{h}\frac{\partial h}{\partial x} \approx \frac{1}{h_0}\frac{\partial h_1}{\partial x}$, $h^3 e^{-\eta h} \frac{\partial h}{\partial x} \approx h_0^3 e^{-\eta h_0} \frac{\partial h_1}{\partial x}$, and $h^3 e^{-2\eta h} \frac{\partial h}{\partial x} \approx h_0^3 e^{-2\eta h_0} \frac{\partial h_1}{\partial x}$. Hence, Eq. (21) linear in $h_1$ is found as

$$\frac{\partial h_1}{\partial t} - \beta h_0^3 \frac{\partial^6 h_1}{\partial x^6} + \alpha h_0^3 \frac{\partial^4 h_1}{\partial x^4} + \frac{3}{h_0}\frac{\partial^2 h_1}{\partial x^2} - \zeta \bar{\sigma} \eta h_0^3 e^{-\eta h_0} \frac{\partial^2 h_1}{\partial x^2} - 2\zeta\eta h_0^3 e^{-2\eta h_0} \frac{\partial^2 h_1}{\partial x^2} = 0\,. \quad (23)$$

As a final step, we substitute $h_1 = |h_1| e^{(iqx+st)}$ into Eq.(23) to find the dispersion relation. Noting that $h_0 = 1$ in dimensionless units

$$s h_1 = -\beta q^6 h_1 - \alpha q^4 h_1 + 3 q^2 h_1 - \eta \zeta e^{-\eta} q^2 h_1 (\bar{\sigma} + 2e^{-\eta})\,,$$

$$\Rightarrow s = q^2(-\beta q^4 - \alpha q^2 + 3 - \eta \zeta e^{-\eta}(\bar{\sigma} + 2e^{-\eta})) \quad (24)$$

In Eq. (24), the terms associated with the spontaneous curvature and the electrostatic repulsion are always negative. Electrostatic interactions stabilize the interface because (i) any undulation locally increases the membrane electrostatic potential, (ii) the crests of the undulated surface experience less repulsion than its troughs. Although the latter effect is negligible when $h \gg \lambda_D$ , the former effect suffices to stabilize the interface in this limit. At even greater heights, electrostatic interactions can be neglected altogether. For a flat membrane with a finite spontaneous curvature $c_0$ , the energy gain e.g. at a crest would be equal to the energy cost at a neighboring trough, thereby eliminating a runaway growth of thermally induced fluctuations. In order for $c_0$ to destabilize the interface, which is observed in bulk vesicles[7], the interface must be curved in the limit $q \to 0$ , as is the case for closed membranes. Thus, the spontaneous-curvature induced destabilization is inherently a non-linear effect.

In contrast, Eq. (24) predicts that attractive van der Waals interactions between the bilayer and the aluminum substrate (i.e., $A_H > 0$), which is known to destabilize a free interface that confines a liquid[6], cause an instability in the bilayer profile. Briefly, any forming crest would be attracted less than the neighboring troughs, resulting in an increasing pressure gradient across, in turn amplifying the fluctuation further. This positive feedback cycle between the bilayer height, liquid pressure, and the resulting flow of the confined liquid leads to an instability of the lipid membrane conformation.

In our experiments, no spontaneous curvature $c_0$ is measured, and the thickness of the lipid bilayer ($\sim 5$ nm) is considerably bigger than the Debye screening length $\lambda_D \approx 0.84$ nm. Then, if we ignore the electrostatic interactions and set $c_0 = 0$, the dimensionless height evolution equation and its dispersion relation Eq. (24) simplify to

$$\frac{\partial h_1}{\partial t} - \beta h_0^3 \frac{\partial^6 h_1}{\partial x^6} + \frac{3}{h_0}\frac{\partial^2 h_1}{\partial x^2} = 0\,, \tag{25}$$

$$s = q^2(-\beta q^4 + 3)\,. \tag{26}$$

The critical wave number $q_c$, which is defined by the relation $s(q_c) = 0$, is calculated as

$$q_c = \left(\frac{3}{\beta}\right)^{1/4} \approx 23.40\,. \tag{27}$$

The critical wave number in real units then yields

$$q_c = \frac{23.40}{L} = 2.34 \times 10^7\ m^{-1}\,. \tag{28}$$

#### S3.5 Numerical procedure and extended results

By only considering the van der Waals interactions, we have solved Eq. (25), which is sixth order in space and first order in time, subject to the six boundary conditions that simulate the ventral region of a bilayer vesicle on a substrate: At $x = 0$, which corresponds to the center of the ventral region, the flow speed $U = 0$ and $\frac{\partial h_1}{\partial x}|_{x=0} = 0$ by symmetry. Furthermore, the membrane curvature must be constant at $x = 0$, leading to $\frac{\partial^3 h_1}{\partial x^3}|_{x=0} = 0$ due to the vanishing flow. At $x = 1$, we take $p = 0$ $\frac{\partial^2 h_1}{\partial x^2}|_{x=0} = 0$ since the the pressure must be balanced by the liquid pressure above the bilayer. We further impose $h_1 = 0$ at $x = 1$. With $h(q, t = 0) = 1 + |h_1|e^{iqx}$, $k_B T \sim 10^{-21}\,J$, $k \sim 10^{-19}\,J$, $q \sim 10^8\,m^{-1}$ and $L \sim 10^{-6}\,m$, we estimate the amplitude of the initial fluctuations using Eq. (4) as $|h_1| = 10^{-11}\,m$. Thus, in dimensionless units, the initial profile of the membrane fluctuations $h_1$ that satisfies the six boundary conditions is given by

$$h_1 = 0.001\cos qx\,, \quad q = (2n+1)\pi/2\,, \quad n = 0,1,2\ldots \tag{29}$$

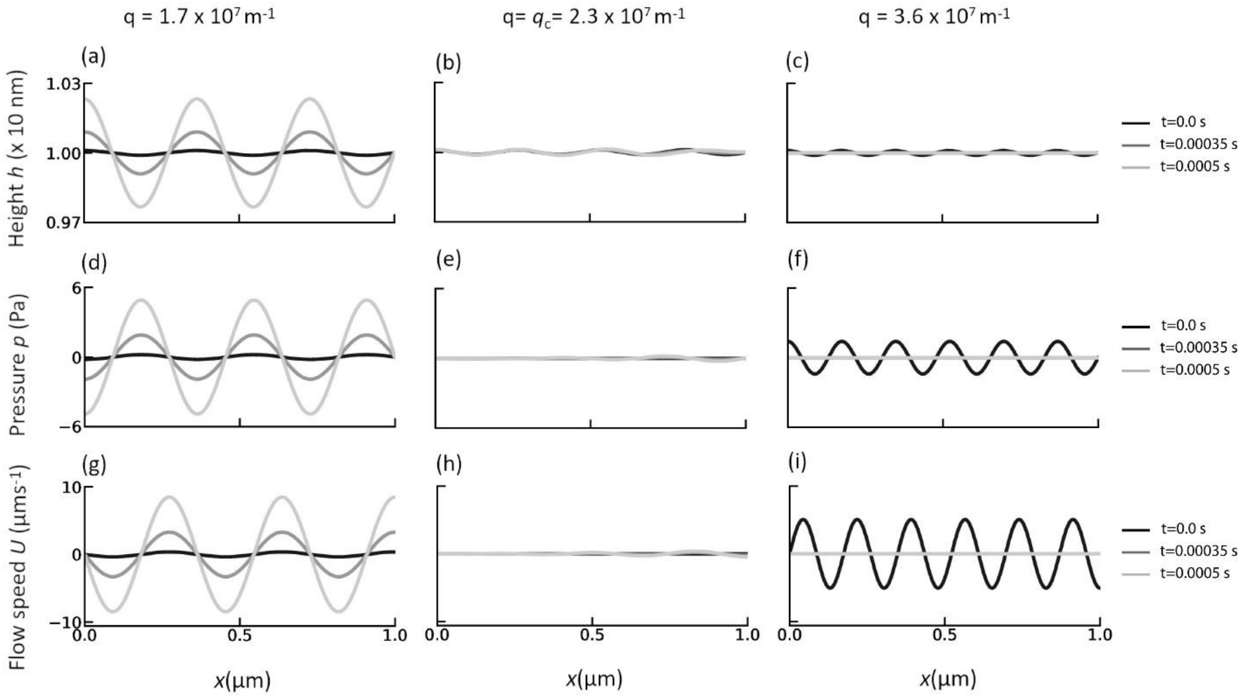

**Figure S3.2: Numerical results in the linear membrane deformation regime.** Height, liquid pressure and flow speed profiles for $q < q_c$, $q = q_c$ and $q > q_c$ at three different times. The center point at the bottom of the vesicle corresponds to $x = 0$, and the contact line is located at $x = 1$ .The grayscale corresponds to three different times $t = 0\ s$, $t = 0.00035\ s$ and $t = 0.0005\ s$. The results from numerical solutions show emergence of the instability below critical wave number $q_c = 2.3 \times 10^7\ m^{-1}$, equivalently above critical wavelength $\lambda_c = \frac{2\pi}{q_c} = 273\ nm$, in agreement with experimentally observed minimum size of the membrane bulges forming at the ventral region of the vesicle.

For numerical implementation we have used the open source finite element analysis library FEniCS on Python 3.6[8]. **Fig. S3.2** shows the time evolution of the height $h$ (panels **a**-**c**), liquid pressure $p$ (panels **d**-**f**) and the height-averaged flow speed $U$ (panels **g**-**i**) for three different wave numbers $q = 1.7 \times 10^7\ m^{-1}$ (left column), $q = q_c = 2.3 \times 10^7\ m^{-1}$ (middle column), and $q = 3.6 \times 10^7\ m^{-1}$ (right column). Below a critical wave number $q < q_c$, the inverse deformation time scale $s$ (Eq. (26)) is positive such that fluctuations grow in time, leading to an instability. Because the troughs of $h$ (**Fig. S3.1**) coincide with the crests of $p$ (**Fig. S3.2d**), the flow profile (**Fig. S3.2g**) resulting from the pressure gradient further amplifies the height undulations. At $q = q_c$, the inverse deformation time scale $s$ is zero, thus, the height profile (**Fig S3.2b**) remains stationary. This steady state is characterized by zero pressure and vanishing flow throughout the system (**Fig. S3.2e** and **h**). For a wave number $q > q_c$, the inverse deformation time scale $s$ is negative, and the fluctuations die out. The maxima and minima of the height profile (**Fig. S3.2c**) coincide, respectively, with the maxima and minima of the pressure profile (**Fig. S3.2f**). Thus, the resulting flow (**Fig. S3.2i**) suppresses the height undulations for all wave numbers $q > q_c$. These results suggest that the height instability triggered by the attractive van der Waals interactions between the lipid bilayer membrane and the aluminum substrate is a plausible mechanism for the birth of vesicular subcompartments in a closed lipid bilayer system.

#### S3.6 Debye-Hückel approximation for electrostatic potential between lipid bilayer and aluminum substrate

This section supplements **S3.2**. The Debye screening length $\lambda_D$ is defined as[3]

$$\lambda_D \equiv \sqrt{\frac{D\epsilon_0 k_B T}{2z^2 e^2 c_\infty}}, \tag{30}$$

where $D$ is the dielectric constant of the ionic solvent, $\epsilon_0$ is the vacuum permittivity, $k_B$ is Boltzmann's constant, $T$ is the temperature, $z$ is the valency of the ions in the solution, $e$ is the charge of an electron and $c_\infty$ is the concentration of the ions far from lipid bilayer. Then, the electrostatic potential $V(z)$ between two charged parallel surfaces in the presence of an aqueous ionic background solvent can be described by the Debye-Hückel equation[3]

$$\frac{d^2V(z)}{dz^2} = \frac{1}{\lambda_D^2} V(z) \tag{31}$$

when the corresponding electrostatic potential energy is much smaller than the thermal energy $k_B T$. The general solution of Eq. (31) is of the form

$$V(z) = Ae^{-z/\lambda_D} + Be^{z/\lambda_D}, \tag{32}$$

where the constants $A$ and $B$ are determined by the boundary conditions. Based on Eq. (31) the potential $V_1(z)$ in the region between aluminum surface and the lipid membrane (subdomain 1) and the potential $V_2(z)$ in the region between lipid membrane and infinity (subdomain 2) are given by

$$V_1(z) = A_1 e^{-z/\lambda_D} + A_2 e^{z/\lambda_D}, \tag{33}$$

$$V_2(z) = B_2 e^{-(z-h)/\lambda_D}. \tag{34}$$

To determine $A_1$, $A_2$ and $B_2$, we must specify three boundary conditions. Denoting the surface charge density of the bilayer by $\sigma_m$, the surface charge density of the aluminum substrate by $\sigma_s$, these three boundary conditions are ($E_1(z) = -\partial V_1/\partial z$, $E_2(z) = -\partial V_2/\partial z$ : the electric fields in subdomains 1 and 2)

$$E_1(0) = \frac{\sigma_s}{D\epsilon_0}, \tag{35}$$

$$E_1(h) + \frac{\sigma_m}{D\epsilon_0} = E_2(h)\,, \tag{36}$$

$$V_1(h) = V_2(h)\,. \tag{37}$$

Substituting Eq. (33) and Eq. (34) in the boundary conditions (35)−(37), we obtain following three equations for the constants $A_1$, $A_2$ and $B_2$ as

$$A_1 = \frac{\lambda_D}{2D\epsilon_0}\left(2\sigma_s + \sigma_m e^{-(h/\lambda_D)}\right)\,, \tag{38}$$

$$A_2 = \frac{\sigma_m \lambda_D}{2D\epsilon_0} e^{-(h/\lambda_D)}\,, \tag{39}$$

$$B_2 = \frac{\sigma_s \lambda_D}{D\epsilon_0} e^{-(h/\lambda_D)} + \frac{\sigma_m \lambda_D}{2D\epsilon_0} e^{-(2h/\lambda_D)} + \frac{\sigma_m \lambda_D}{2D\epsilon_0}\,. \tag{40}$$

The solutions to the electrostatic potential $V(z)$ in the subdomains 1 and 2 are then given by

$$V_1(z) = \frac{\lambda_D}{2D\epsilon_0}\left[2\sigma_s + \sigma_m e^{-h/\lambda_D}\right] e^{-z/\lambda_D} + \frac{\sigma_m \lambda_D}{2D\epsilon_0} e^{(z-h)/\lambda_D}\,, \tag{41}$$

$$V_2(z) = \left[\frac{\sigma_s \lambda_D}{D\epsilon_0} e^{-(h/\lambda_D)} + \frac{\sigma_m \lambda_D}{2D\epsilon_0} e^{-(2h/\lambda_D)} + \frac{\sigma_m \lambda_D}{2D\epsilon_0}\right] e^{-(z-h)/\lambda_D}\,. \tag{42}$$

Furthermore, the electric field profile is given as

$$E_1(z) = -\frac{\partial V_1}{\partial z} = \frac{1}{2D\epsilon_0}\left(2\sigma_s + \sigma_m e^{-(h/\lambda_D)}\right) e^{-z/\lambda_D} - \frac{\sigma_m}{2D\epsilon_0} e^{(z-h)/\lambda_D}\,, \tag{43}$$

$$E_2(z) = -\frac{\partial V_2}{\partial z} = \frac{\sigma_s}{D\epsilon_0} e^{-z/\lambda_D} + \frac{\sigma_m}{2D\epsilon_0} e^{-(z+h)/\lambda_D} + \frac{\sigma_m}{2D\epsilon_0} e^{-(z-h)/\lambda_D}\,. \tag{44}$$

The electrostatic potential and the electric field profiles are shown in **Fig. S3.3**. Therefore, the electric field on the bilayer is governed by the combined effect of the substrate and the screening charges in between, i.e.,

$$E(h) = \lim_{z \to h^-} E_1(z) = \frac{\sigma_s}{D\epsilon_0} e^{-h/\lambda_D} + \frac{\sigma_m}{2D\epsilon_0}\left(e^{-2h/\lambda_D} - 1\right)\,, \tag{45}$$

for which the potential at $z = h$ becomes

$$V(h) = \frac{\lambda_D \sigma_s}{D\epsilon_0} e^{-h/\lambda_D} + \frac{\sigma_m \lambda_D}{2D\epsilon_0} e^{-2h/\lambda_D} + \frac{\sigma_m \lambda_D}{2D\epsilon_0}\,. \tag{46}$$

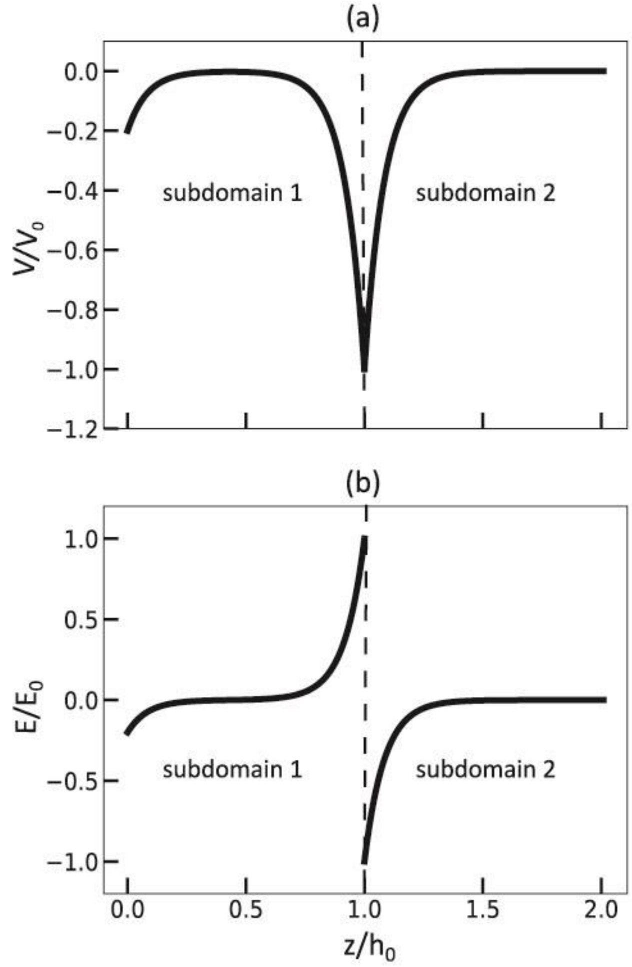

**Figure S3.3: The electrostatic potential and the electric field** in the subdomain 1 ($0 \leq z/h_0 \leq 1$) and subdomain 2 ($1 < z/h_0$). The electrostatic potential (**a**) is continuous at the subdomain boundary while the electric field (**b**) is discontinuous due to surface charge present on the lipid bilayer. Here $V_0 = \frac{|\sigma_m|\lambda_D}{2D\epsilon_0}$ and $E_0 = \frac{|\sigma_m|}{2D\epsilon_0}$.

### S4. Fluorescein encapsulation

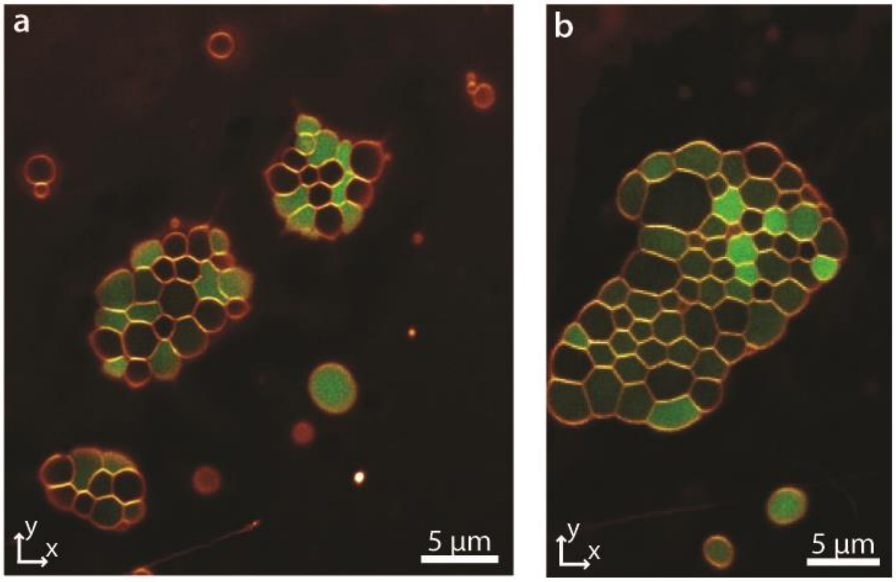

**Figure S4.** Encapsulation of fluorescein. (**a-b**) Confocal micrographs of protocell superstructures after exposure to the fluorescein (top view). Subcompartments encapsulate fluorescein (green) at different concentrations.

### S5. DNA encapsulation

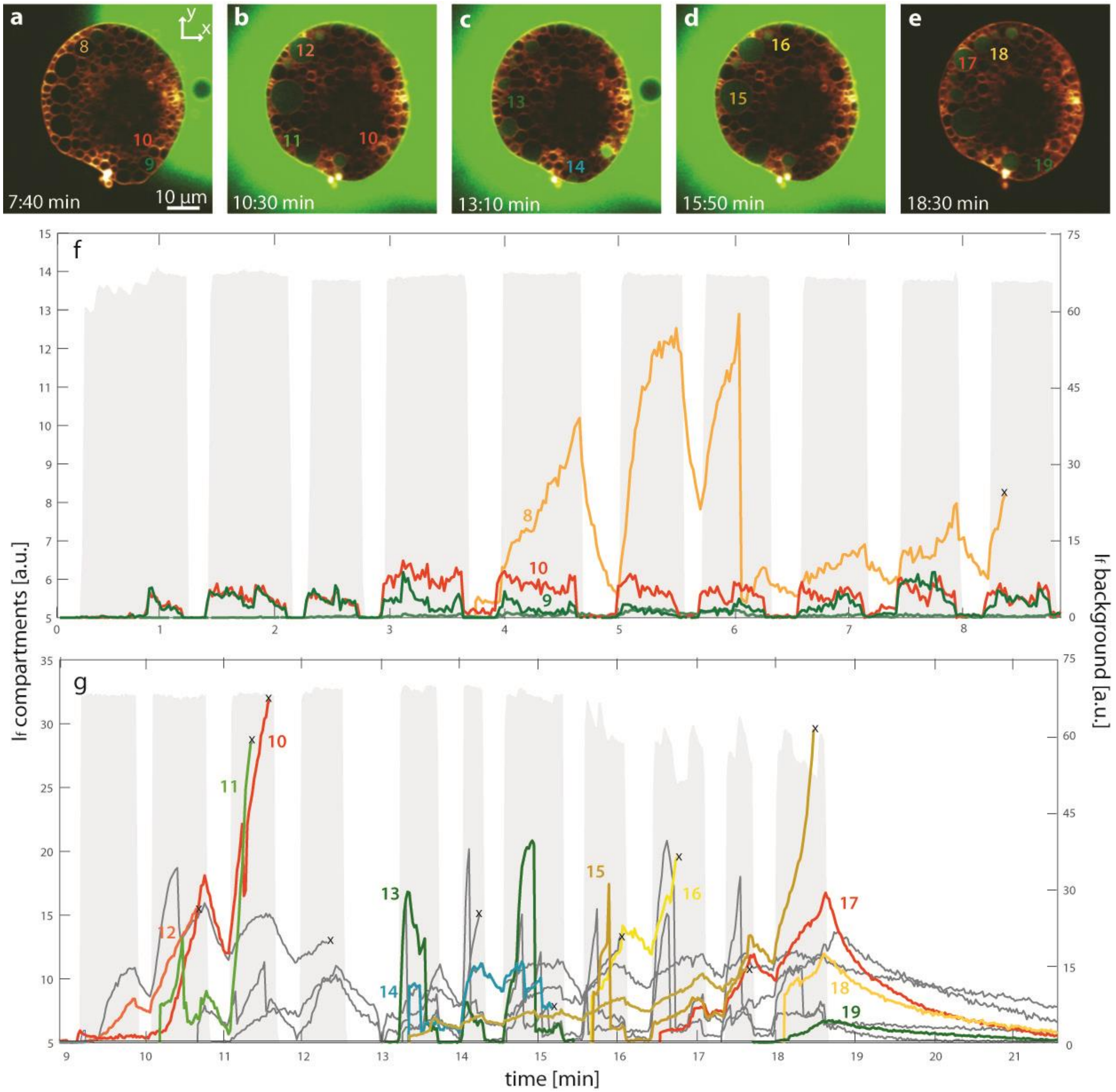

**Figure S5.** Full graph of FAM-DNA encapsulation experiment shown in **Fig. 5a-i**. (**a-e**) Confocal micrographs showing the fluorescently labeled DNA encapsulated inside protocell superstructure in different time points. (**f-g**) Graph showing DNA fluorescence intensity versus time. For clarity, the graph of a continuous experiment is divided into two parts: 0-9 min (**a**) and 9-21.5 min (**b**). Colored plots show intensity in 12 individual subcompartments, each plot corresponds to one subcompartment in **a-e** labeled with the same number. Dark grey plots correspond to the plots depicted in Fig. 4i, grey zones depict intensity of the recirculation zone in the exterior region of the superstructures. The plots marked with 'x' show subcompartments which could not be monitored further.

### Supporting Movies

**Movie S1. Growth and dynamics of protocell superstructures.** Laser scanning confocal microscopy time series showing temperature-induced emergence, growth and unification of protocell colonies.

**Movie S2. Mechanical stability of protocell superstructures.** Laser scanning confocal microscopy time series showing the mechanical stability of individual model protocells and protocell superstructures exposed to osmotic pressure.

**Movie S3. DNA encapsulation.** Laser scanning confocal microscopy time series showing encapsulation of fluorescently labeled DNA inside superstructures and exo-compartments.

**Movie S4. DNA strand displacement reaction.** Laser scanning confocal microscopy time series showing DNA strand displacement reaction inside superstructures and pseudo-division.

### References

1. Tu, Z. C.; Ou-Yang, Z. C., Lipid membranes with free edges. *Physical Review E* **2003,** *68* (6), 061915.
2. Zhong-can, O.-Y.; Helfrich, W., Bending energy of vesicle membranes: General expressions for the first, second, and third variation of the shape energy and applications to spheres and cylinders. *Physical Review A* **1989,** *39* (10), 5280-5288.
3. Phillips, R.; Kondev, J.; Theriot, J.; Garcia, H. G.; Orme, N., *Physical biology of the cell*. 2013.
4. Israelachvili, J. N., 6 - Van der Waals Forces. In *Intermolecular and Surface Forces (Third Edition)*, Israelachvili, J. N., Ed. Academic Press: San Diego, 2011; pp 107-132.
5. Doi, M., *Soft matter physics*. First edition. ed.; Oxford University Press: Oxford ;, 2013.
6. Oron, A.; Davis, S. H.; Bankoff, S. G., Long-scale evolution of thin liquid films. *Reviews of Modern Physics* **1997,** *69* (3), 931-980.
7. Steinkühler, J.; Knorr, R. L.; Zhao, Z.; Bhatia, T.; Bartelt, S. M.; Wegner, S.; Dimova, R.; Lipowsky, R., Controlled division of cell-sized vesicles by low densities of membrane-bound proteins. *Nature Communications* **2020,** *11* (1), 905.
8. Alnæs, M. S.; Blechta, J.; Hake, J.; Johansson, A.; Kehlet, B.; Logg, A.; Richardson, C.; Ring, J.; Rognes, M. E.; Wells, G. N., The FEniCS project version 1.5. *Archive of Numerical Software* **2015,** (Vol. 3, No. 100).
